# The mechanism of selective kinesin inhibition by kinesin binding protein

**DOI:** 10.1101/2020.07.17.208736

**Authors:** Joseph Atherton, Jessica J. A. Hummel, Natacha Olieric, Julia Locke, Alejandro Peña, Steven S. Rosenfeld, Michel O. Steinmetz, Casper C. Hoogenraad, Carolyn A. Moores

## Abstract

Subcellular compartmentalisation is necessary for eukaryotic cell function. Spatial and temporal regulation of kinesin activity is essential for building these local environments via control of intracellular cargo distribution. Kinesin binding protein (KBP) interacts with a subset of kinesins via their motor domains, inhibits their microtubule (MT) attachment and blocks their cellular function. However, its mechanisms of inhibition and selectivity have been unclear. Here we use cryo-electron microscopy to reveal the structure of KBP and of a KBP-kinesin motor domain complex. KBP is a TPR-containing, crescent-shaped right-handed α-solenoid that sequesters the tubulin-binding surface of the kinesin motor domain, structurally distorting the motor domain and sterically blocking MT attachment. KBP uses its α-solenoid concave face and edge loops to bind the kinesin motor domain and selective mutation of this extended binding surface disrupts KBP inhibition of kinesin transport in cells. The KBP-interacting surface of the motor domain contains motifs exclusively conserved in KBP-interacting kinesins, providing a basis for kinesin selectivity.

## Introduction

Kinesins are a superfamily of microtubule (MT)-based molecular motors that play important roles in cellular functions such as mitosis, cell motility and intracellular transport^1-3^. Kinesins are categorised into 14 sub-classes (kinesin-1 to kinesin-14^4^) by motor domain conservation and within these sub-classes individual family members (a total of 45 ‘Kif’ genes in humans and mice) have a wide range of functional characteristics and biological roles^1,5^. Dysfunction of kinesin family members has been implicated in a number of pathological conditions^6,7^. The kinesin motor domain is the microtubule-binding engine that drives these activities, converting the chemical energy of ATP binding and hydrolysis into mechanical force. While these mechanical forces are classically used to generate motility in transport kinesins, some kinesin family members drive MT organisation or depolymerisation of MTs.

Kinesins are highly regulated in order to prevent both waste of ATP and to spatially and temporally control kinesin function. This is particularly important in highly polarised and compartmentalised cells such as neurons. Kinesin regulation via inhibition of their motor domains can occur through a number of mechanisms that limit ATPase activity and/or block track binding - these include intramolecular inhibition by kinesin tail domains, post-translational modification of the motor, or through interactions with regulatory binding partners. Recently, it has been demonstrated that a subset of kinesin superfamily members, including kinesin-2s, −3s, −8s and −12s, are sequestered by kinesin binding protein (KBP; KIF1BP; KIAA1279), which inhibits MT track attachment by their motor domains and, thus, blocks their MT-related functions^8-10^.

KBP is expressed in multiple human tissues including brain and heart^8^. Mutations in the KBP have been identified as causing autosomal recessive Goldberg-Shprintzen syndrome (GOSHS)^11-14^, which presents as congenital facial dysmorphia, nervous system pathology and dysfunction and heart defects^15^. In addition, KBP gene copy number has been recently reported as predictive in paediatric neuroblastoma prognosis, prompting its suggestion as a drug target^16^. KBP was originally identified as a kinesin-3 binding partner that modulated its mitochondrial transport function^8^; however KBP has since been shown to interact with a subset of other kinesin family members to regulate diverse cellular processes including mitosis^17,18^, spermatogenesis^19^ and neuronal differentiation, growth and cargo distribution^10,20-24^.

We do not currently know what the structure of KBP is, nor understand the mechanism of KBP-kinesin inhibition. It is also completely unknown how KBP differentiates between particular kinesin family members. KBP is a 72kDa protein, is predicted to contain several tetratricopeptide repeats (TPRs) and to be mainly α-helical in secondary structure content^8,9^. Here we present cryo-electron microscopy (cryo-EM) structures of KBP alone and of KBP bound to the motor domain of the mitotic kinesin Kif15 (a 110kDa complex). We show that KBP is a TPR-containing right-handed α-solenoid protein composed of 9 antiparallel α-helix pairs interrupted by a linker region. We also show that KBP’s concave face binds Kif15 via its MT-binding elements and induces a large displacement of the kinesin α4 helix, sterically inhibiting MT association. Finally, we show that KBPs kinesin selectivity is associated with specific kinesin sequences spread across the interaction surface.

## Results

### KBP is a TPR-containing right-handed α-solenoid

The 3D structure of the ∼72kDa KBP at 4.6 Å resolution (Fig. 1 and Supplementary Fig. 1a,b) was determined using cryo-EM data collected using a Volta Phase Plate (VPP), and an atomic model was calculated (see Methods). Our structure revealed that KBP is a right-handed α-solenoid protein (Fig. 1a,b and Supplementary Fig. 1c-e). Nine pairs of anti-parallel α-helices (αHP1 (α-helical pair 1) to αHP9) are broken by a single ‘linker α-helix’ (LαH) and ‘linker loop’ (LL) in the centre of the fold separating KBP into N-terminal and C-terminal subdomains (Fig. 1 and Supplementary Fig. 1c-f). The four predicted TPR motifs contribute exclusively to α-helical pairs in the N-terminal subdomain (Fig. 1a,d,e).

**Figure 1.**
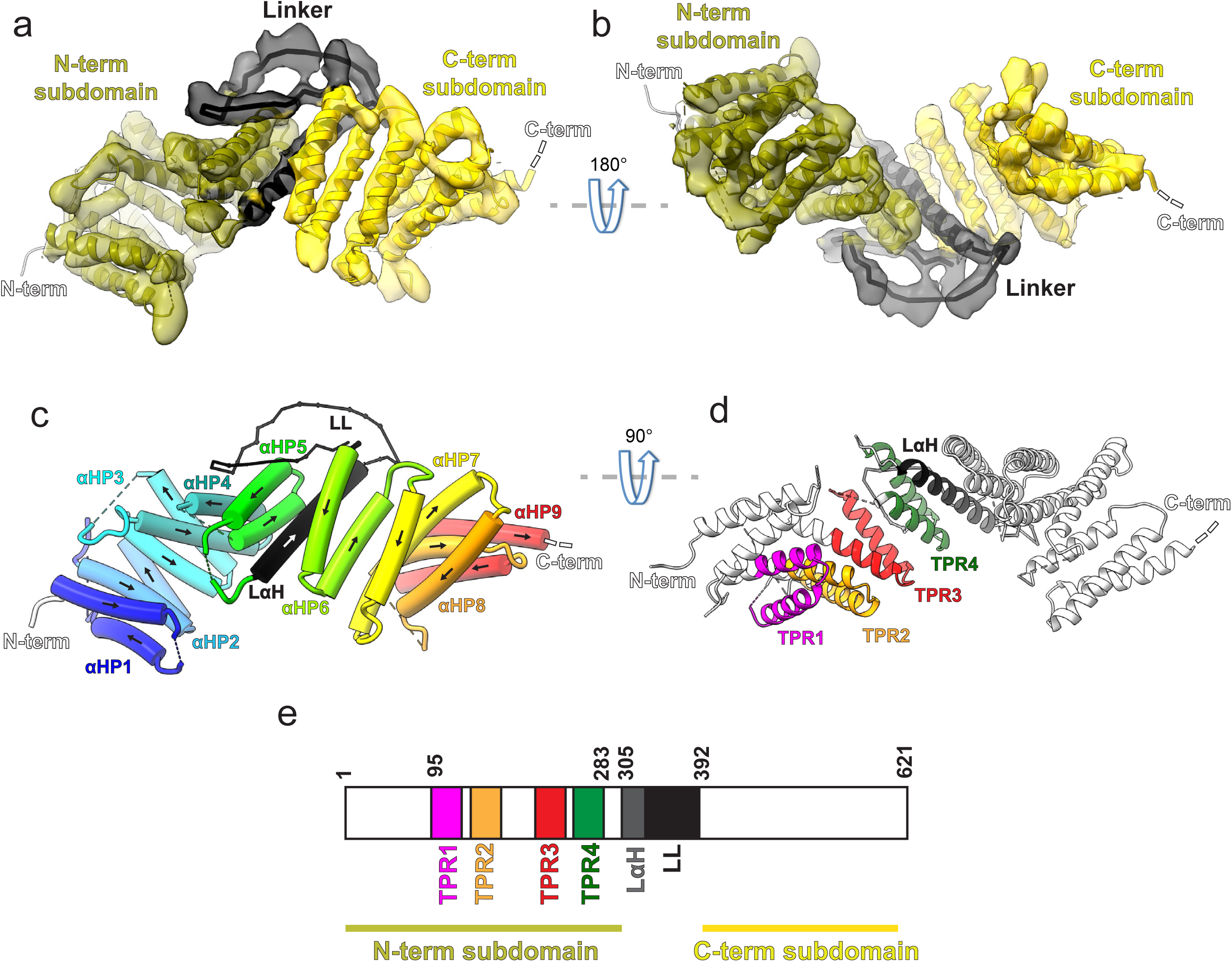
KBP is a TPR-containing right-handed α-solenoid. (a) Model of KBP (ribbon representation) displayed in experimental cryo-EM density. The N-terminal (olive) and C-terminal (gold) subdomains are separated by a linker region (black). Semi-transparent density is coloured regionally as per the fitted model. The N- and C-termini are shown, with a dotted line representing the disordered C-terminus (not modelled). The linker loop region was not modelled but its rough path is indicated as a solid black line. (b) The same as panel a, but rotated 180° around the axis indicated. (c) The same view as in panel a, but with the density removed and α-helices displayed as pipes with their directionality indicated by arrows. The 9 antiparallel α-helical pairs (αHP1 to αHP9) are each coloured separately and labelled, as is the linker α-helix (LαH) and linker loop (LL). (d) Ribbon representation of KBP showing the 4 tetratrico peptide repeat (TPR) motifs and the LαH coloured according to the labels. View related to panel c, by a 90° rotation around the indicated axis. (e) Schematic of the KBP showing the position of the TPR motifs between residue 95 and 283 of the N-terminal subdomain and position of the linker region (LαH and LL) between residues 305 and 392.

The supercoiling α-helical pairs form concave and convex faces linked by short and long loops that constitute the two edges of the α-solenoid (Supplementary Fig. 1c-f). In contrast to the shorter loops, the longer loops (>7 residues) tend to be partially disordered, show low sequence homology between KBPs in different species and are mainly found in the N-terminal subdomain (e.g. L2, L6 and L10, Supplementary Fig. 1d-f). The linker loop is the longest (62 residues) and is thus unique in the KBP structure because it is reasonably conserved and mainly ordered, with visible corresponding density clearly bridging the N and C-terminal subdomains (Fig. 1a-b and Supplementary Fig. 1d-f). Despite this clear ordered density, this loop was not modelled due to low homology to available structures and a lack of consensus in secondary structure prediction (see Methods). In spite of this lack of consensus, density in this region suggests that part of this loop may form further α-helical structures. Other TPR-containing α-solenoid proteins form important regulatory interactions in numerous contexts, and the structure we describe is indicative of similar properties for KBP.

### KBP conformationally adapts to bind Kif15’s motor domain using both subdomains

To elucidate the mechanism of kinesin inhibition by KBP, we determined the structure of KBP in complex with the Kif15 (kinesin-12) motor domain (Kif15_MD). The overall resolution of this KBP-Kif15_MD complex was 6.9 Å, with KBP and Kif15_MD determined to similar local resolutions (Supplementary Fig. 2a-b). We built a model of the complex via flexible fitting using our KBP model and the Kif15_MD crystal structure (Fig. 2a,b and see Methods). The complex is arranged such that Kif15_MD sits in the concave face of the KBP α-solenoid, analogous to a baseball enclosed in a baseball glove. The kinesin MD is positioned centrally between the N and C-terminal subdomains and contacts the KBP concave face and loops at the α-solenoid edges.

**Figure 2.**
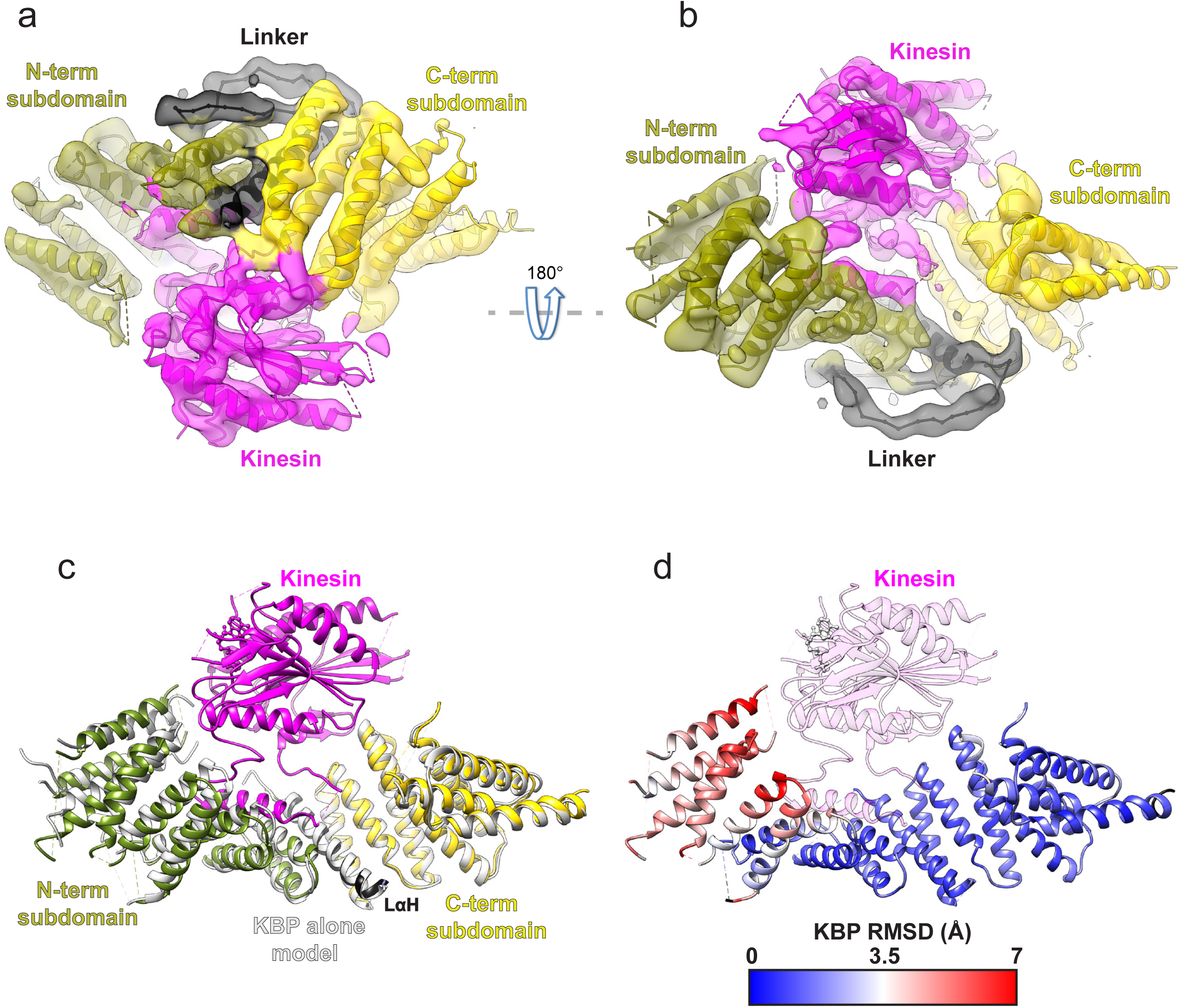
KBP conformationally adapts to bind Kif15’s MD via both subdomains. (a) Model of the KBP-Kif15_MD complex (ribbon representation) displayed in experimental cryo-EM density. The N-terminal (olive) and C-terminal (gold) subdomains and the linker region (black) are shown in KBP, while kinesin is coloured in magenta. Semi-transparent density is coloured regionally as per the fitted model. (b) The same as panel a, but rotated 180° around the axis indicated. (c) The KBP-alone model (light grey ribbons) was superimposed on the KBP-Kif15_MD model (opaque ribbons) using Chimera’s matchmaker^56^. Colouring and view as in panel b. (d) RMSD in Å for KBP comparing KBP-Kif15_MD and superimposed KBP-alone models as in panel c, shown on KBP from the KBP-Kif15 model. Parts of the KBP model coloured black are disordered/missing in the KBP alone model. The Kif15_MD is shown in transparent magenta.

When the structure of KBP-alone is superimposed onto KBP in the KBP-Kif15_MD complex, it is clear that KBP undergoes a conformational change in the presence of its kinesin binding partner, with the largest differences resulting from an unfurling motion of the N-terminal subdomain (Fig. 2c,d and Supplementary Video 1). The KBP-alone model is incompatible with Kif15_MD binding, due to clashes with L14 in the C-terminal subdomain and αHP3a, αHP4a and L8 in the N-terminal subdomain. The conformational changes in KBP upon Kif15_MD binding relieves these clashes in the complex (Supplementary Video 1).

To establish whether the KBP-Kif15_MD mode of interaction applied to other kinesins, we also collected data of the complex formed by KBP with the motor domain of the kinesin-3 Kif1A (Kif1A_MD). 2D classification of these images revealed a number of classes with an extra-density the size of a kinesin motor domain bound to the concave face of KBP, consistent with what was observed in the KBP-Kif15_MD dataset (Supplementary Fig. 2c). However, in contrast to the KBP-Kif15_MD sample, these KBP-Kif1A_MD 2D classes provided only limited views of the complex (Supplementary Fig. 2c,d), such that a reliable 3D structure could not be calculated. Intriguingly, in addition, the extra kinesin density in the 2D classes appeared to have a somewhat flexible position relative to KBP. However, these data did allow us to confirm that indeed Kif1A_MD also interacts with KBP on its concave face in the same way as Kif15_MD and suggests a common mechanism of kinesin inhibition by KBP.

### Kif15_MD binds KBP via rearrangement of its tubulin-binding subdomain

We examined the effect of KBP binding on the conformation of Kif15_MD. Kinesin motor domains can be structurally divided into three distinct subdomains^25,26^ which undergo coordinated conformational changes during the MT-based kinesin ATPase cycle. MT binding stabilises the tubulin-binding subdomain while the P-loop and Switch 1/2 subdomains – which contain the conserved nucleotide-coordinating P-loop and Switch 1 and 2 motifs - move relative to each other in response to the nucleotide state of the MD^25-27^. We determined the structure of the MT-bound, AMPPNP state of Kif15_MD, which shows that this MD adopts a canonical conformation (Supplementary Fig. 3). Comparison of this conformation with an ADP-bound Kif15_MD crystal structure (PDB: 4BN2^28^) illustrates the scale of these MT- and nucleotide-dependent subdomain rearrangements in Kif15, which are similar to those seen in other kinesins MDs^25,27,29^ (Supplementary Fig. 3d,e and Fig. 3c,d).

**Figure 3.**
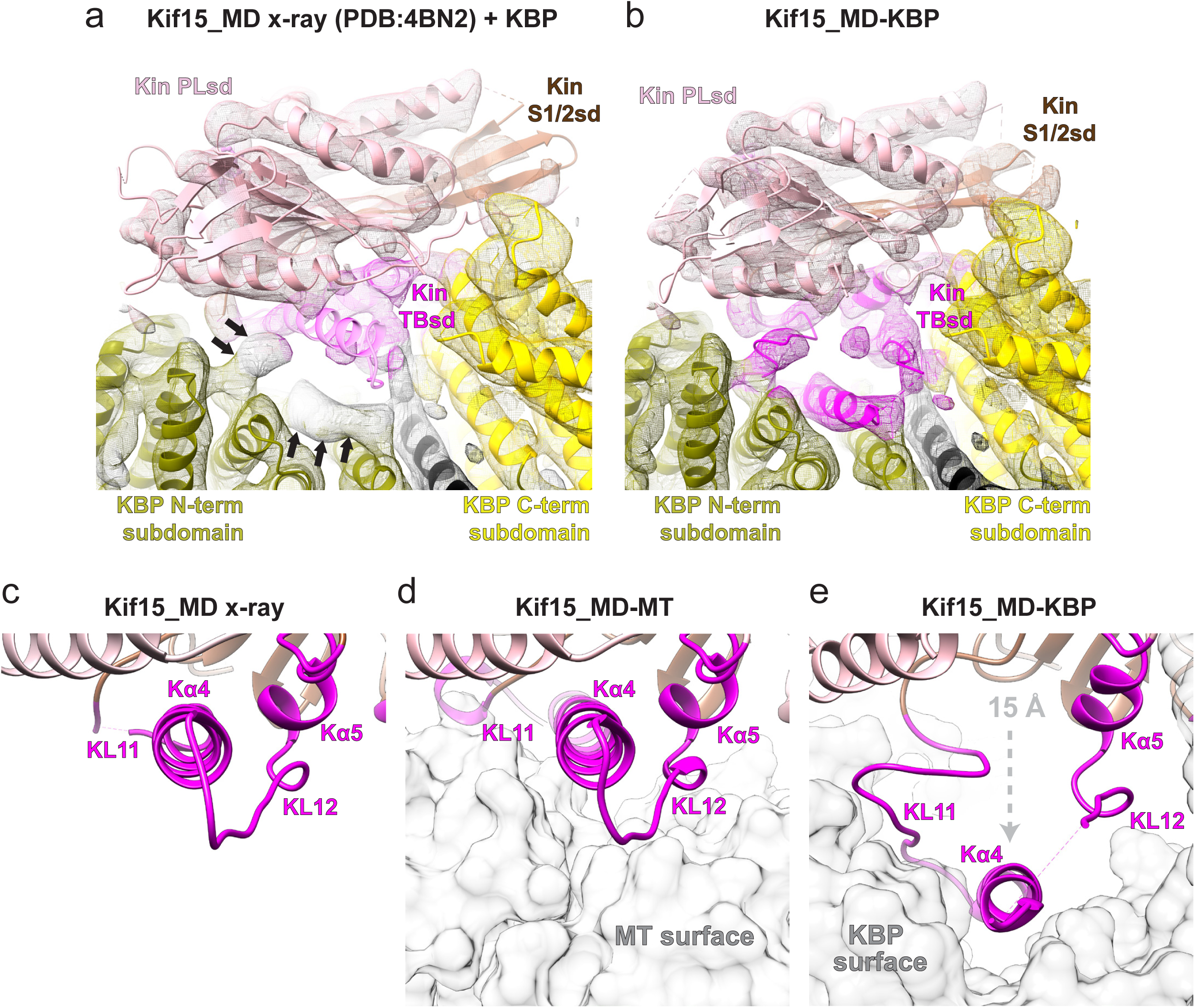
Kif15_MD binds KBP via rearrangement of its tubulin-binding subdomain. (a) The crystallographic model of the Kif15_MD alone (PDB: 4BN2^28^) was superimposed on the Kif15_MD region of the KBP-Kif15_MD complex, with the Kif15_MD of the KBP-Kif15_MD complex model hidden. The Kif15_MD Switch 1/2 subdomain (Switch 1/2 subdomain) is coloured sienna, the P-loop subdomain (Kin-PLsd) is coloured light pink. The TBsd of the Kif15_MD crystallographic model is shown as pale magenta to illustrate poor fit into density. The KBP subdomains are coloured as labelled. Black arrows indicate unaccounted-for cryo-EM density. Individual secondary structure elements in the tubulin-binding subdomain are labelled. The cryo-EM density for the KBP-Kif15_MD complex is shown in mesh and is coloured by proximity (≤ 3.5 Å) to the fitted model. (b) Same as in panel a, but the whole fitted KBP-Kif15_MD complex model is shown. The Kif15_MD tubulin-binding subdomain (TBsd) is now coloured magenta to indicate good fit into density. (c) Zoomed view of just the TBsd (corresponding to the boxed region in Supplementary Fig. 4d), showing just the Kif15_MD-alone crystallographic model. (d) The TBsd in the Kif15_MD-MT model, same view as in panel c. The MT is shown in light grey surface representation. (e) The TBsd in the KBP-Kif15_MD-complex model, same view as in panel c. KBP is shown in light grey surface representation and the ∼15 Å displacement of helix α4 is indicated by the dashed grey arrow.

The structure of the KBP-Kif15_MD complex revealed that KBP binds the kinesin motor domain via the tubulin-binding subdomain (Fig. 3). While the P-loop and Switch 1/2 subdomains of the Kif15_MD crystal structure and associated Mg_2+_-ADP generally fitted well into density of the KBP-Kif15_MD complex, a large portion of the tubulin-binding subdomain did not (Fig. 3a; Supplementary Fig. 4a,b). In particular, there is a striking lack of density in the expected position for helix α4 (Fig. 3a; Supplementary Fig. 4b). Instead, there was strong density of length and width consistent with helix α4 displaced by ∼15 Å into the concave face of KBP, and which we modelled as such (Fig. 3b,e, Fig. 2 and Supplementary Fig. 4b-e). This displacement of helix α4, which lies close to the TPR-repeat region of the N-terminal subdomain of KBP, is accompanied by additional rearrangements of the flanking L11 and L12 in Kif15_MD (labelled KL11 and KL12, Fig. 3c-e: Supplementary Fig. 4b-e). A number of other TPR-containing α-solenoids are known to bind peptide motifs with α-helical content within their concave faces^30-32^ (Supplementary Fig. 4f-i), and our structure shows that KBP binds helix α4 of Kif15_MD in a similar way.

The KBP-bound conformation of the Kif15_MD tubulin-binding subdomain is also radically different from its MT-bound conformation (Fig. 3d-e). The tubulin-binding subdomain forms the majority of the MT-binding surface in the Kif15_MD-MT complex (Fig. 3d, Supplementary Fig. 3) such that KBP and MTs cannot simultaneously bind Kif15_MD due to extensive steric overlap (Fig. 3d-e). In summary, KBP sequesters and blocks the MT-interacting surface of kinesin motor domains via a mechanism that involves significant conformational change within the motor domain.

### KBP binds kinesin motor domains via conserved motifs in the α-solenoid edge loops and α-helices at the concave face

KBP contacts the Kif15_MD both via 1) loops connecting the α-solenoid edges and 2) TPR-containing α-helices at the concave face (Fig. 4, Supplementary Fig. 5 and Supplementary Video 2). At the α-solenoid edges, L1, L3, L5 and L10 in the N-terminal subdomain and L12, L14, L16 and L18 in the C-terminal subdomain are close enough to Kif15_MD to be involved in binding. The closest interaction of these were KBP L12 and L14, which contact both Kβ5-KL8 and KL12-Kα5-KL13 regions of the Kif15 tubulin-binding subdomain (Fig. 4). KBP’s disordered L1 lies close to Kif15_MD’s KL9, while the shorter, ordered L3 and L5 are situated near but not contacting KL11 and Kα6 (Supplementary Fig. 5a,b). KBP’s C-terminal L16 and L18 are close enough to Kif15_MD that they may interact with the flexible KL12, N-terminus or neck-linker. At the TPR-containing region of the concave face of KBP, αHP4a, αHP4b and αHP5a contact the K11-Kα4-KL12 region of Kif15_MD (Fig. 4c-d).

**Figure 4.**
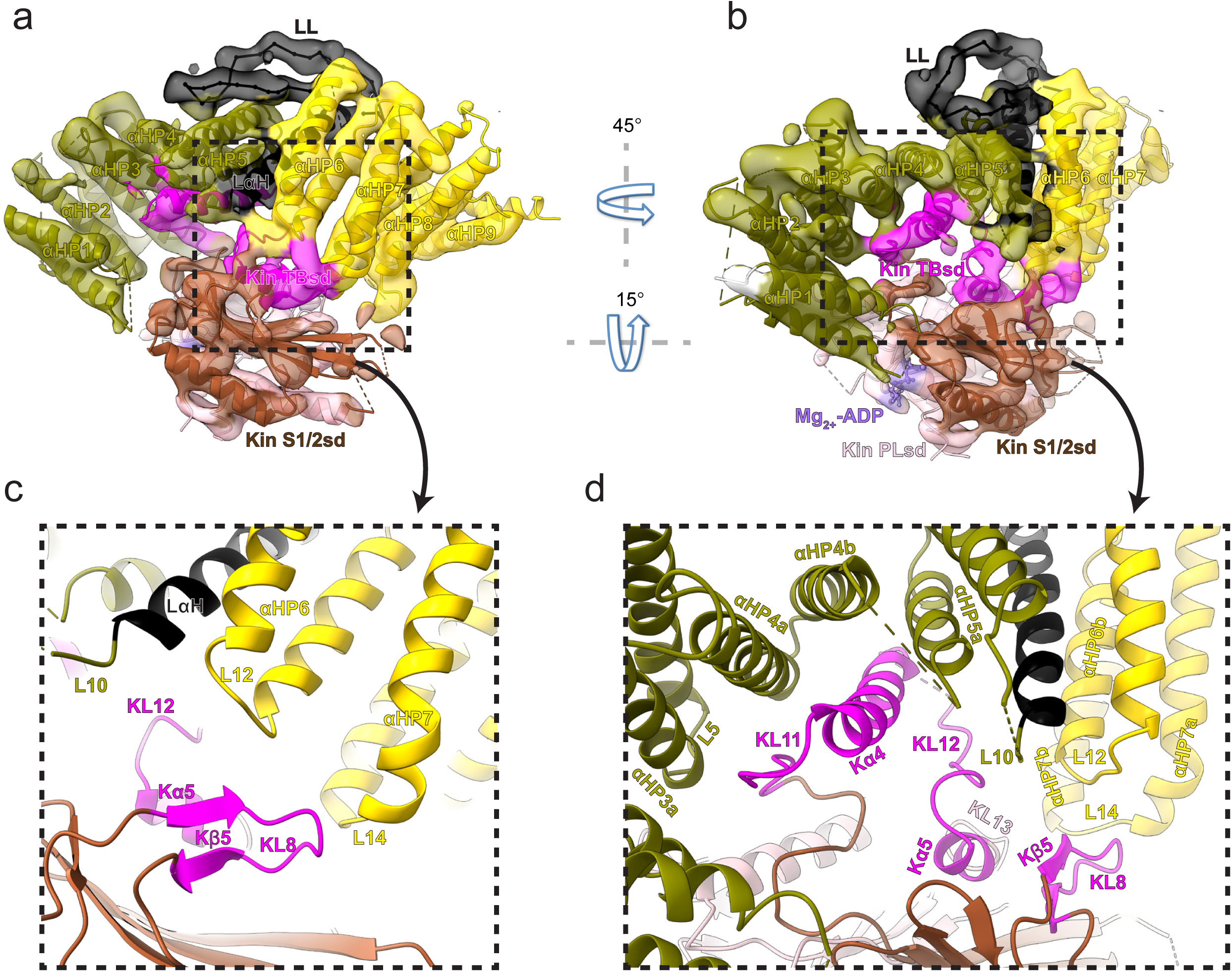
KBP binds kinesin MDs via conserved motifs in the α-solenoid edge loops and α-helices at the concave face. (a) Pseudo-atomic model of the KBP-Kif15_MD complex (ribbon representation) displayed in cryo-EM density, using the same viewpoint as Fig. 2a, but with the Kif15_MD now coloured by subdomain as in Fig. 3. The Kif15_MD Switch 1/2 subdomain (Switch 1/2 subdomain) is coloured sienna, the P-loop subdomain (Kin-PLsd) is coloured light pink. The Kif15_MD tubulin-binding subdomain (TBsd) is coloured magenta. The KBP subdomains are coloured as labelled. The nine helix pairs of KBP are labelled. Semi-transparent density is coloured regionally as per the fitted model. (b) The same as panel a, but rotated 45° and 15° respectively around the axes indicated. (c) Zoomed view of the region indicated in panel a, with density removed and selected Kif15_MD and KBP secondary structure elements labelled. (d) Zoomed view of the region indicated in panel b, with density removed and selected Kif15_MD and KBP secondary structure elements labelled.

To test the functional significance of this interface, we investigated KBP-kinesin interactions in cells. First, we expressed in COS7 cells wild-type or mutant KBP constructs, in which the predicted interacting amino acids within the loops of interest were substituted for Ala, Gly or Pro residues (Supplementary Fig. 1f and Supplementary Table 2), and assessed inhibition of kinesin motility using inducible peroxisome transport assays (Fig. 5a-c). In this assay, dimeric Kif15 or Kif1A constructs consisting of only the motor domain, the first coiled-coil region and an FRB-tag (Kif15_MDC-FRB or Kif1A_MDC-FRB) are expressed together with PEX-mRFP-FKBP, a peroxisome-binding construct, and with various KBP constructs. Addition of rapalog induces FRB-FKBP heterodimerisation and motor-driven peroxisome transport into the cell periphery, except when the motor is inhibited by KBP (Fig. 5a-c). Second, we also used pull-down assays to assess the motor-KBP interaction. For this, Kif1A_MDC or Kif15_MDC was fused to bioGFP and co-expressed with HA-tagged mutant KBP constructs in HEK293T cells, followed by pull-down of HA-KBP constructs by the bioGFP-Kif_MDC^9^ (Supplementary Fig. 6).

**Figure 5.**
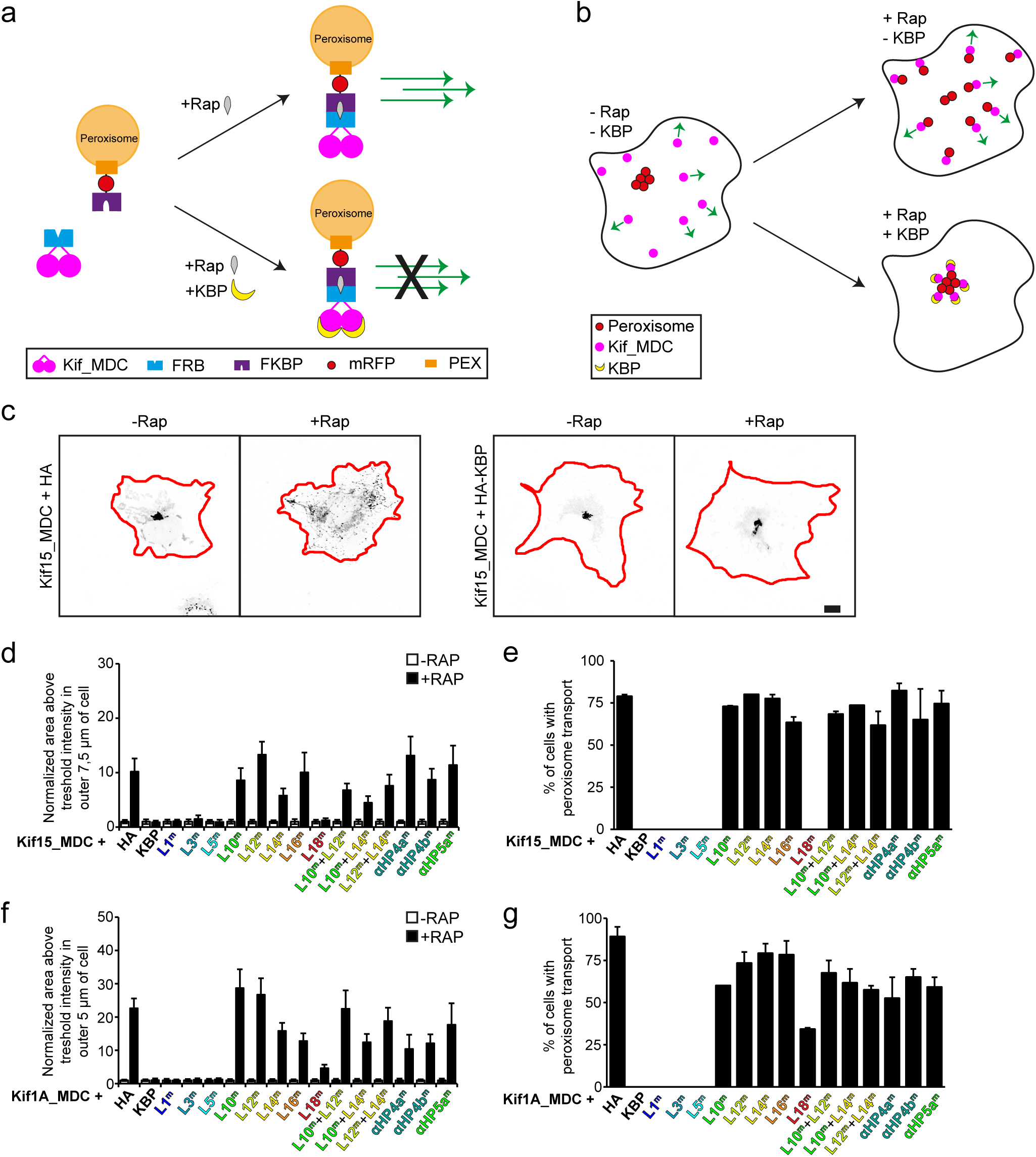
Disruption of cryo-EM defined KBP-kinesin interface perturbs KBP inhibition of Kif15 and Kif1A motility in cells. (a) Schematic depiction of the inducible peroxisome motility assay, with the kinesin motor domain fused to an FRB domain and PEX fused to an FKBP domain. Addition of rapalog links FRB and FKBP and induces peroxisome movement by kinesin dimers. Expression of KBP inhibits kinesin movement, such that addition of rapalog cannot induce peroxisome transport. (b) Schematic representation of the inducible peroxisome motility assay in cells. Without rapalog or KBP, peroxisomes localize in the cell center, whereas kinesin moves towards the cell periphery. Rapalog induces peroxisome transport into the cell periphery, which is inhibited in presence of KBP. (c) Representative images of peroxisomes in COS7 cells expressing Kif15_MDC-FRB, PEX-mRFP-FKBP and HA (left panels) or HA-KBP (right panels) without and with addition of rapalog. Scale bar, 10 μm. (d, f) Quantification of the area above threshold intensity in the outer 5 μm (Kif1A_MDC) or 7.5 μm (Kif15_MDC) of the cell in cells expressing Kif15_MDC-FRB (d) or Kif1A_MDC-FRB (f), PEX-mRFP-FKBP, and HA-KBP constructs without and with rapalog treatment. Values are normalized to the condition without rapalog. Data are displayed as mean ± s.e.m. (n=28-35 cells from two independent experiments). (e, g) Quantification of the percentage of cells in which peroxisome movement is observed after rapalog treatment in cells expressing Kif15_MDC-FRB (e) or Kif1A_MDC-FRB (g), PEX-mRFP-FKBP, and HA-KBP constructs. Data are displayed as mean ± s.e.m. (n=28-35 cells from two independent experiments).

Consistent with our structural analysis, mutation of either of L12 or L14 reduced KBP interaction with Kif15_MDC/Kif1A_MDC by pull-down assay (Supplementary Figure 6b,c); mutation of L12 or L14 also strongly abrogated KBP inhibition of induced Kif15_MDC/Kif1A_MDC-based transport, as can be seen by an increase of peroxisomes in the cell periphery after rapalog addition similar to the control condition without KBP (Fig. 5d-g,). Mutation of the 20-residue long L10 had a similar effect, suggesting some part of this loop may interact with either α5 or KL12 of Kif15/Kif1A (Fig. 5d-g, Supplementary Fig. 6b,c). The above described mutations were also combined to assess additive effects. Here we observed that KBP constructs containing mutations in both L10 + L12, L10 + L14 or L12 + L14 had similar effects to KBP with only one of the loops mutated in both the inducible peroxisome transport assay and pull-down assay (Fig. 5d-g, Supplementary Fig. 6b-c).

Although mainly disordered in the structure, mutation of the C-terminal 16 residue long L16 also disrupted KBP inhibition of Kif1A_MDC and Kif15_MDC (Fig. 5d-g, Supplementary Fig. 6b,c). L16 contains a number of conserved residues and may flexibly interact with the core of the P-loop subdomain and Switch 1/2 subdomain or the disordered regions of the kinesin N-terminus, neck-linker or KL12 (Supplementary Fig. 1f & Supplementary Fig. 6b,c). Interestingly, mutation of L18 reduced KBP interaction with Kif1A_MDC but not Kif15_MDC as assessed by pull-down assay or inducible peroxisome transport assay. In contrast, mutation of L1, L3 or L5 had no effect on KBP’s interaction with either Kif15_MDC or Kif1A_MDC, (Fig. 5d-g, Supplementary Fig. 6b,c) suggesting that these elements do not significantly contribute to complex formation.

Ala-substitutions in the TPR-containing α-helices at the KBP concave face were targeted at particularly inter-species conserved polar residues predicted to interact with the Kif15_MD K11-Kα4-KL12 region in the reconstruction (Fig. 4c,d and Supplementary Fig. 1f). Ala substitution of KBP Tyr-213 and Gln-216 in αHP4a, or Gln-238 in αHP4b strongly reduced KBP’s interaction with both Kif15_MDC and Kif1A_MDC in pull-down assays and also reduced KBP’s ability to inhibit peroxisome transport via these motors (Fig. 5d-g, Supplementary Fig. 6b,c). Ala-substitution of Thr-255 and Gln-258 in αHP5a had similar effects.

In summary, while the various tested mutations reduced KBP-Kif15_MDC and KBP-Kif1A_MDC binding, none completely blocked the interaction, including the mutations that targeted two of the loops from L10, L12 and L14. These mutation studies thus support the idea that KBP interacts with different kinesin family members in a similar way via an extended interface at KBP’s concave face that is composed of TPR-containing α-helices and α-solenoid edge loops.

### Specific sequences in the tubulin-binding subdomain are conserved across KBP-binding family members

Given that KBP selectively binds and inhibits only a subset of kinesins^9^, we used our structural data to investigate the basis of this selectivity. Analysis of the tubulin-binding subdomain sequences in KBP-binding and KBP-non-binding kinesins (Fig. 6a) revealed patterns of sequence conservation across the entire subdomain in all KBP-binding kinesins; this included both the Kβ5-KL8 and KL11-Kα4-KL12-Kα5-KL13 regions. In contrast, the equivalent regions are variable in kinesins that are not inhibited by KBP. The length of KL8, which joins the two Kβ5 strands, was also consistently 5 residues long in KBP-binding kinesins, while it was variable in KBP-non-binding kinesins. From our KBP-Kif15_MD structure, the sensitivity of the KBP interaction to KL8 length makes sense considering the tight fit of this loop between KBP L12 and L14 (Fig. 6b). In summary, two consensus motifs in KL11-Kα4-KL12-Kα5-KL13 and Kβ5-KL8 regions of the tubulin-binding subdomain are found exclusively in KBP-binding kinesins and these are likely to form the basis of KBP’s kinesin family member selectivity. We therefore propose a model where KBP selects and inhibits target kinesins through binding and remodelling a compatible tubulin-binding subdomain, obstructing the kinesin MT-binding surface (Fig. 6c).

**Figure 6.**
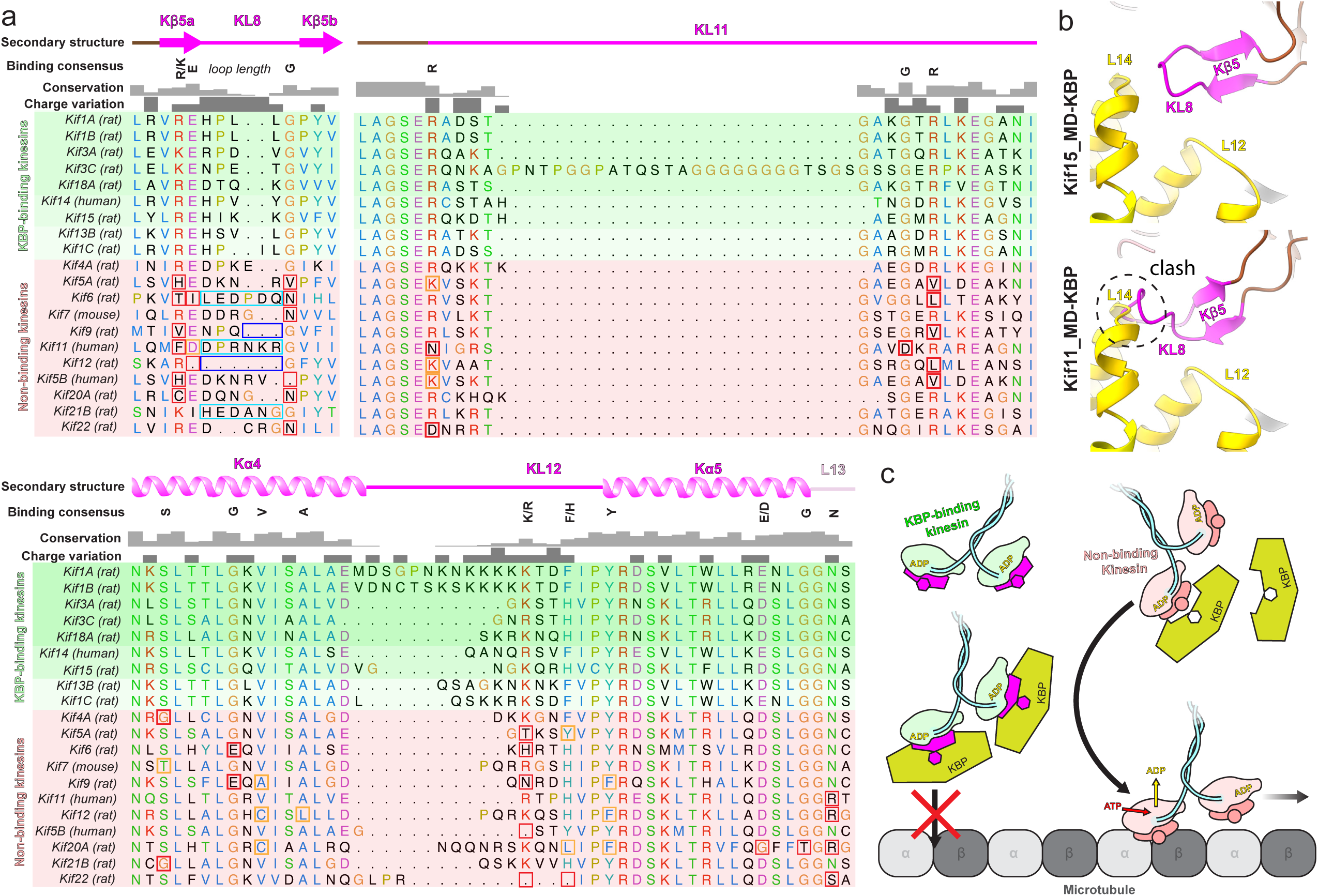
Conserved motifs in KBP-binding kinesin MDs. (a) Sequence alignment of the tubulin-binding subdomain from kinesin motor domains, made using Clustal Omega multiple sequence alignment^66^. Residues are coloured according to standard Clustal X colouring (dependent on residue type and conservation, see http://bioinfolab.unl.edu/emlab/documents/clustalx_doc/clustalx.html#C). Kinesin MD constructs experimentally assessed for KBP interactivity are taken from^9^; strongest interactors are in rows highlighted in darker shades of green, weaker interactors in lighter shades of green and non-interactors in red. Secondary structure element, conservation and charge variation columns, as well as a ‘binding consensus’ column indicating residues/loop length conserved at the interface (according to the Kif15_MD-KBP complex) in KBP-binding but not non-binding kinesins are shown above the alignment. Non-conservation relative to this consensus is shown in boxed sequence; red boxes, non-conservative substitutions, orange boxes, conservative substitutions (general charge/polarity/hydrophobicity retention), cyan boxes, extended loop region, dark blue boxes, truncated loop region. (b) Top panel; view of the Kβ4-KL8 region of the tubulin-binding subdomain in the KBP-Kif15_MD model, coloured as in Fig. 4. Bottom panel; as in upper panel, but with the Kif11_MD cryo-EM model (PDB: 6TA4^75^) superimposed onto the now hidden KBP-Kif15_MD. Note steric clash introduced by Kif11_MD’s extended KL8. c) Schematic model of KBP’s selective kinesin inhibition mechanism. KBP (olive) binds the compatible TBsd of recognised kinesins (magenta) but is incompatible with the TBsd of non-binding kinesins (salmon). For its target kinesins, KBP therefore sterically blocks the TBsd interaction with MTs (grey), preventing activation of kinesin ATPase and motility.

## Discussion

In this study, we reveal the TPR-containing right-handed α-solenoid structure of the ∼72kDa KBP using VPP cryo-EM. At the time of writing and to our knowledge, structures of only a few macromolecular complexes <80 kDa have been determined using cryo-EM^33-37^. Further, the structure of the KBP-Kif15_MD complex shows how KBP binds the Kif15_MD via its concave face and undergoes subtle remodelling of its N-terminal domain to accommodate kinesin binding. This further reinforces the idea that the TPR-containing structures are not simply static scaffolds but can flexibly respond to ligand binding^38^. In contrast, the Kif15 motor domain undergoes a radical conformational change in forming a complex with KBP, in which helix α4, the major component of the motor’s tubulin-binding subdomain, is displaced from the main MD body by ∼15 Å into the KBP concave face. This is consistent with this region of the kinesin motor domain being rather malleable and able to move independently of the core structure of kinesin motor domains^39,40^. This displacement expands the surface area over which the normally compact tubulin-binding subdomain can interact with KBP, and it is through the sequence and shape of this interface that the selectivity of KBP for a subset of kinesin motors is defined.

Analysis of the KBP-Kif1A_MD complex (Supplementary Fig. 2c) supports the idea of a conserved mode of interaction between a subset of kinesins and the concave face of KBP. Interestingly, Kif1A_MD exhibited flexibility in its interaction with KBP, which was not observed in the KBP-Kif15_MD complex. Whether this reflects a physiological reality, or a result of the EM preparation method is uncertain at present. However, our 2D classifications combined with mutation studies strongly suggest Kif15_MD and Kif1A_MD share an overall similar KBP binding mode. In fact, the only notable difference in the mutation analysis was Kif1A’s distinctive sensitivity to mutation of KBP’s L18, which may derive from its proximity to Kif1A’s kinesin-3 family-specific flexible basic KL12 extension (the K-loop).

Targeted mutations to the various kinesin-binding KBP elements reduced complex affinity, yet no single mutation completely disrupted the interaction, possibly explaining why only nonsense mutations/deletion of the KBP gene have been reported in GOSHS^11-14^. Our structure shows that KBP binds exclusively to the tubulin-binding subdomain of Kif15_MD, sterically blocking the kinesin tubulin binding surface, preventing MT attachment. Interaction of kinesin motor domains with the MT surface via the tubulin-binding subdomain stimulates nucleotide exchange and kinesin ATPase activity. However, the displacement of helix α4 from the kinesin nucleotide-binding site, together with the absence of MT-mediated ordering of KL9 and KL11, means that while interaction with KBP stabilises the kinesin tubulin-binding subdomain, its catalytic site is distorted and the structural changes associated with MT-stimulated ATPase cannot occur. This could be an important facet of the role of KBP in the energy economy of the cell in addition to directly blocking kinesin-MT interactions.

The concave face of TPR-containing α-solenoids commonly serve as a recognition platform for specific peptide motifs, including those forming α-helical structures^41^. Specificity and affinity for target motifs are determined in part by the shape of the α-solenoid concave face, which in turn is defined by the fold supertwist. In addition, particular amino-acid arrangements at the concave face contribute to partner binding affinity and specificity, together with additional interfaces formed at the convex surface or α-solenoid edge^42^. KBP kinesin specificity and affinity is defined by the interaction of its concave face with the large surface area of the kinesin L11-Kα4-KL12-Kα5-KL13 region, in addition to binding the kinesin Kβ5-KL8 region at its α-solenoid edge. Interestingly, distal to the N-terminal MT-binding region of kinesin-1, C-terminally-associated kinesin light chain use their unique TPR-containing α-solenoid concave face to select cargos via recognition of specific peptide motifs^43^. Therefore, peptide selectivity by TPR-containing α-solenoids is a facet of both kinesin MT-binding and cargo-binding regulatory mechanisms. Such protein-protein interactions may be selectively targeted for disruption^44^, and the insights arising from our work provide future avenues to disrupt KBP-kinesin interactions and thereby explore KBP interactions and regulatory roles.

The effective and selective kinesin inhibitory mechanism of KBP revealed by our work may fulfil specific roles in the kinesin regulatory toolbox employed by cells to spatially and temporarily orchestrate kinesin activity. Future studies will be aimed at understanding how KBP interacts with kinesins in dimeric and/or autoinhibited forms. It will also be of key importance to elucidate the mechanisms of KBP activity regulation, for example by phosphorylation of KBP and/or kinesins^9^ and KBP acetylation and targeted degradation by the ubiquitin system^45^, and to expand our understanding KBP’s biological roles in neuronal function and cancer.

## Methods

### Protein expression and purification for cryo-EM

Full length human KBP residues 1-621 in a PSTCm1 expression vector (with kanamycin resistance and a N-terminal thrombin cleavable 6 x His-tag) was expressed in Rosetta2 cells (Novagen). Following immobilised metal-affinity chromatography with Ni-NTA resin (Qiagen), the 6 x His-tag was removed via incubation with thrombin protease overnight at 4 °C. The protein was then subjected to reverse IMAC and further purified using size exclusion chromatography (SEC) into a buffer of 20mM TrisHCL (pH7.4), 150mM NaCl, 2.5mM CaCl2, 1mM DTT. Protein was snap-frozen and stored in at −80 °C.

A human Kif15 motor domain and neck linker construct (Kif15_MD residues 1-375) in a pET21a vector with a C-terminal 6 x His-tag was generated by chemical synthesis (GenScript, Piscataway, NJ). Six of the eight cysteine residues (C5S, C50S, C162S, C294S, C314S, C346S) were mutated and two cysteines were inserted (S250C, G375C) for orthogonal experiments not described further here. The construct was expressed and purified using methods previously described^46^, then buffer exchanged into 25 mM HEPES pH 7.5, 100 mM KCl, 2 mM MgCl2, 1 mM EGTA, 1 mM DTT, 0.1 mM ATP for storage.

A human Kif1A motor domain and neck linker construct (Kif1A_MD residues 1-362) in a pFN18a vector (with a TEV protease-cleavable N-terminal Halo-tag and a C-terminal 6 x His-tag) was expressed in BL21-Gold (DE3) cells. Following a first IMAC with Ni-NTA resin, the Halo-tag was removed via incubation overnight with TEV protease at 4 °C. The protein was then isolated from TEV via a second IMAC with Ni-NTA resin and further purified by SEC into a storage buffer of (20 mM HEPES, pH 7, 150 mM NaCl, 5 mM MgCl_2_, 0.1 mM ADP and 1 mM TCEP).

Kif15_MD or Kif1A_MD complexes with KBP were purified via IMAC using the 6 x His tag on the kinesin constructs. Briefly, His-tagged kinesins were incubated with a 10 times excess of KBP in 20 mM TrisHCL (pH7.5), 150 mM NaCl, 1 mM MgCl_2,_ 10 mM Imidazole, 1 mM DTT, 0.2 mM ADP for 5 minutes at 4 °C. Following IMAC, complexes were eluted from the Ni-NTA resin (Qiagen) by addition of 200 mM imidazole, then dialyzed at 4 °C for 4 hours into 20 mM TrisHCL (pH7.5), 150 mM NaCl, 1 mM MgCl_2_, 1 mM DTT, 0.2 mM ADP.

### Sample preparation for cryo-EM

KBP was prepared for cryo-EM using three different approaches. In the first approach, KBP was diluted to 0.15 mg/ml in KBP dilution buffer (20 mM TrisHCL, pH7.5, 150 mM NaCl, 2 mM DTT) and 4 μl were applied to glow-discharged C-flat™ 2/2 holey carbon EM grids (Protochips, Morrisville, NC). For the second approach, KBP was diluted to 0.3 mg/ml in KBP dilution buffer and 4 μl were applied to glow-discharged 1.2/1.3 AuFoil gold grids (Quantifoil^®^). For the third approach, glow-discharged 1.2/1.3 AuFoil gold grids (Quantifoil^®^) were coated with graphene-oxide (GO) according to the protocol described by Cheng and colleagues^47^ then 4 μl of KBP diluted to 0.02 mg/ml in KBP dilution buffer were added.

Kinesin motor domain-KBP complexes were diluted to 0.03 mg/ml in KBP-kinesin dilution buffer (20 mM TrisHCL (pH7.5), 50 mM NaCl, 1 mM MgCl_2_, 1 mM DTT, 0.2 mM ADP) and 4 μl were added to the GO-coated gold grids described above. After a 30 second incubation of samples on the EM grid in a Vitrobot Mark IV (FEI Co., Hillsboro, OR) set at 4 °C and 80 % humidity, samples were blotted (6-8 seconds, blot force −10) and vitrified in liquid ethane. All steps were performed at 4 °C.

For preparation of the Kif15_MD-MT complex, porcine tubulin (>99% pure, Cytoskeleton Inc.) was polymerised in MES polymerisation buffer (100 mM MES, 1 mM MgCl_2_, 1 mM EGTA, 1 mM DTT, pH 6.5) with 5 mM GTP at 37 °C then stabilised with 1 mM paclitaxel. ∼70 μM Kif15_MD was pre-incubated for 5 min with 5 mM of AMPPNP in BRB80 at room temperature, and then mixed with 20 μM stabilised MTs. After a further incubation of 15 min, a 4 μl droplet was applied to a pre-glow discharged holey carbon grid (2/2 C-flat, Protochips Inc.), blotted for 3.5 s and then vitrified in liquid ethane using a Vitrobot Mark IV at ambient temperature and 80% humidity.

### Cryo-EM data collection

For dataset of KBP alone or KBP-Kif15_MD, low-dose movies were collected automatically using EPU software (Thermo Fisher, MA, USA) on a Titan Krios electron microscope (Thermo Fisher) operating at 300 kV, with a K2 summit direct electron detector (Gatan, CA, USA) and a quantum post-column energy filter (Gatan) operated in zero-loss imaging mode.

Datasets of KBP alone were collected either at eBIC or the ISMB, Birkbeck using a Volta phase plate (VPP), a sampling of ∼1.05 Å/pixel and a nominal defocus range of 0.5-0.7 μm. The total dose was 42 e-/Å^2^ over 40 frames, with the detector operating in counting mode at a rate of ∼5 e-/pixel/second.

Datasets of KBP-kinesin complexes were collected at the ISMB, Birkbeck without a phase plate and a nominal defocus range of 1.5-4 μm. Kif1A_MD-KBP complexes were collected at a sampling of 0.85 Å/pixel, whereas Kif15_MD-KBP complexes were collected at a sampling of 1.047 Å/pixel. For Kif1A_MD-KBP complexes, the total dose was 88 e-/Å^2^ over 36 frames, with the detector operating in counting mode at a rate of 7.1 e-/pixel/second. For Kif15_MD-KBP complexes, the total dose was 80 e-/Å^2^ over 64 frames, with the detector operating in counting mode at a rate of 5.7 e-/pixel/second.

The Kif15-MT dataset was collected manually on a Tecnai Polara microscope (Thermo Fisher) at the ISMB, Birkbeck, operating at 300 kV, with a K2 summit direct electron detector (Gatan, CA, USA) and a quantum post-column energy filter (Gatan) operated in zero-loss imaging mode. A nominal defocus range of 1.0-3.5 μm and a final pixel size of 1.39 Å was used. The total dose was 32 e-/Å^2^ over 50 frames, with the detector operating in counting mode at a rate of 6.2 e-/pixel/second.

### Cryo-EM data processing

Low-dose movies were motion-corrected using MotionCor2^48^ with a patch size of 5, generating full-dose and dose-weighted sums. CTF determination was performed on full-dose sums with gCTF^49^ and then dose-weighted sums were used for all further processing. Data were cleaned at this stage by first excluding all micrographs with gCTF resolutions worse than 4.5 Å, as estimated with a custom cross-correlation coefficient cutoff (Python script kindly shared by Radostin Danev), then manually removing micrographs with poor appearance (ice contamination, protein aggregation or poor GO coverage) in real or reciprocal space. For KBP alone data, micrographs with calculated phase shifts outside the expected phase shift progression at each plate position were also excluded.

Particles were first picked using Eman2’s neural network picker^50^, with a 180 pixel box size for KBP-alone and KBP-Kif15_MD datasets, or a 220 pixel box size for the KBP-Kif1A_MD datasets. Good 2D classes were then used as templates to pick the data with Gautomatch (http://www.mrc-lmb.cam.ac.uk/kzhang/).

For Eman2 neural network picker or Gautomatch-derived particles from each dataset, separate multiple rounds of 2D classification were performed in RELION v3.0^51^, cryoSPARC2^52^ or cisTEM^53^. This resulted in a total of six sets of good 2D classes showing clear secondary structure for each dataset, two produced by each programme for each picking method. For each dataset, these six good 2D class sets for each dataset were then combined and duplicate particles removed. At this stage, for each sample (KBP-alone, Kif15_MD-KBP or Kif1A_MD-KBP) good 2D classes from their constituent datasets were combined.

Kif15_MD-KBP or Kif1A_MD-KBP datasets, composed of their respective constituent datasets were easily combined, being from the same microscope and optical set up. However, KBP-alone data were collected on different microscopes and had a range of pixel sizes (<2% difference). KBP-alone data therefore was combined at this stage using the optics grouping protocol in RELION v3.1^51^.

KBP-alone and KBP-Kif15_MD data were taken to 3D processing at this stage, while multiple attempts to process KBP-Kif1A_MD data in 3D gave no reliable results. For KBP-alone and KBP-Kif15_MD data, *de novo* initial 3D models were created in cryoSPARC2. For KBP-Kif15_MD data, a single round of 3D classification was performed in RELION v3.0 and the best class selected and auto-refined. For KBP-alone data, 3D classification in RELION v3.1 or cryoSPARC2 did not reveal different 3D structures or improve reconstructions over sorting only in 2D; therefore, particles selected with 2D classification were used as direct input for auto-refinement. The final KBP-alone map was sharpened with a B-factor of −200 to the gold-standard FSC 0.143 cutoff (4.6 Å). The KBP-Kif15_MD map was sharpened locally with a B-factor of −495, according to local resolutions determined using RELION v3.1’s inbuilt local resolution software.

The Kif15_MD-MT dataset was processed using our MT RELION-based pipeline (MiRP) as described previously, using low-pass filtered Kif5B_MD-decorated MTs as references ^54,55^. Kif15_MD 13-protofilament-MTs were the most common MT architecture and were selected after supervised 3D classification in MiRP for analysis. The symmetrised asymmetric unit (Kif15_MD plus a tubulin dimer) was locally sharpened in UCSF Chimera with a B-factor of −134 according to local resolutions determined using RELION v3.0’s inbuilt local resolution software.

All displayed 3D molecular representations were made in UCSF Chimera or ChimeraX software^56,57^. Data collection and model refinement statistics can be found in Supplementary Table 1..

### Cryo-EM model building and refinement

Due to low overall homology to available structures in the protein data bank (PDB), structure prediction of KBP produced poor models with little resemblance to the cryo-EM density. KBP was therefore modelled using a combination of secondary structure prediction, TPR prediction, fragment homology information, prior knowledge of right-handed alpha-solenoid proteins and with reference to the cryo-EM density.

TPR motifs were identified in the KBP sequence using the TPRpred server^58^ available in the MPI Bioinformatics Toolkit^59^. Secondary structure predictions using Raptor X^60^, iTasser^61^, JPred^62^, Spider2^63^, PSSpred^64^ and SOPMA^65^ were then run on the sequence and consensus between these multiple predictions used to assign likely α-helical content. To identify regions dispensable for the overall fold and likely disordered loop regions, disorder prediction was performed with Raptor X and inter-species low homology regions in KBP (from early Metazoans to humans) were determined via Clustal Omega multiple sequence alignment^66^. Finally, weak homology models for overlapping fragments of the structure were identified using the HHpred^67^ server in the MPI Bioinformatics Toolkit.

With the information described above a sequence alignment was built with KBP and the following fragment homology model PDBs; 5OJ8, 4A1S, 3QC1, 4NQ0, 4AIF, and 5MX5. This sequence alignment was used as a basis for multiple rounds of modelling and flexible fitting with Modeller^68^ and Flex-EM^69^ respectively, using α-helical secondary structure restraints. This modelling process was guided by consistency with the cryo-EM density and secondary structure and TPR predictions described above. Finally, the structure was refined against the cryo-EM density in real-space with 5 macro-cycles in Phenix^70^. All 19 predicted and modelled helices were accounted for by rod-like cryo-EM density in the reconstruction and at 4.6 Å resolution, density was discernible for bulky side chains in the TPR regions (Supplementary Fig. 1a,b), providing a validation of the assigned sequence directionality in the fold.

The KBP-Kif15_MD model was built as follows: the final KBP model described above and the Kif15_MD x-ray crystallographic model (PDB code:4BN2^28^) were rigid fitted into the KBP-Kif15_MD density in Chimera. Density for an extended α6-helix and docked neck-linker in Kif15_MD was absent; therefore, Modeller was used to model a short α6-helix and the neck-linker removed. A model for the L11, α4-helix and L12 region in Kif15_MD were then created using Coot and Modeller. The model was refined into the cryo-EM density in real-space using Phenix^7^ with secondary structure restraints. A first refinement of 15 cycles used rigid bodies describing α-helical hairpins in KBP to get a good rough fit. Following this, the whole complex was further refined without rigid bodies for another 5 macro-cycles.

The Kif15_MD-MT model was built as follows: the Kif15_MD, Kif11_MD and Kif5B_MD-tubulin x-ray crystallographic models (PDB codes:4BN2^26,28,71^) were used as homology models in Modeller to build the Kif15_MD part of the complex. The Kif15_MD model and the taxol-MT tubulin dimer model^72^ were then rigid fitted into Kif15_MD-MT density, combined then refined in real-space with 5 macro-cycles in Phenix with peptide backbone restraints.

### Antibodies, reagents and expression constructs for cell biology

The following antibodies were used for immunofluorescence staining: mouse anti-HA (1:500, Roche), and goat-anti-mouse Alexa 488 (1:400, Thermo Fisher Scientific). The following antibodies were used for western blot: mouse anti-HA (1:2,000, BioLegend), rabbit anti-GFP (1:10,000, Abcam), goat anti-mouse IRDye800CW (1:15.000, LI-COR), and goat-anti-rabbit IRDye680LT (1:20,000, LI-COR). A reagent used in this study is rapalog (AP21967, TaKaRa).

The following DNA expression constructs in this study have been described before: GW1-PEX3-mRFP-FKBP1, βactin-KIF1A_MDC-FRB^9^, BirA coding vector^73^ and pebioGFP^73^. pGW1-HA-KBP contained a linker (GGATCCCCGGAATTCGGCACGAGGGAGGCCGCT) between the HA tag and KBP and was cloned using PCR based strategies with human KBP cDNA (KIAA1279, IMAGE clone 4550085) as template and ligation into the pGW1-HA backbone. A similar strategy was used to generate the mutated KBP constructs, listed in Supplementary Table 2. βactin-KIF15_MDC-FRB was cloned using a PCR based Gibson Assembly strategy with mouse KIF15 cDNA as template into the βactin-KIF1A_MDC-FRB backbone. PebioGFP-KIF1A_MDC and pebioGFP-KIF15_MDC were cloned into the pebioGFP backbone using PCR based strategies with MDC-FRB constructs as templates.

### Cell culture, transfection and immunofluorescence staining

COS-7 cells were cultured in 50/50 DMEM (Lonza)/Ham’s F10 (Lonza) medium supplemented with 10% FCS (Sigma) and 1% penicillin/streptomycin (Sigma). One day before transfection cells were diluted and plated on 18-mm glass coverslips. COS-7 cells were transfected using FuGENE6 (Roche) following the manufacturer’s protocol. Next day, rapalog (final concentration 1 μM) was added and cells were incubated for 3 hours. Cells were then fixed with 4% formaldehyde/4% sucrose in phosphate-buffered saline (PBS) for 10 minutes at room temperature, washed three times PBS-CM (PBS supplemented with 1 mM MgCl_2_ and 0.1 mM CaCl_2_), permeabilized in 0.2% TritonX-100 for 15 minutes and washed one time with PBS-CM. Cells were first incubated with 0.2% gelatin for 30 minutes at 37 °C, and then with primary antibodies, diluted in 0.2% gelatin, for 30 minutes at 37 °C. After washing three times with PBS-CM, cells were incubated for 30 minutes at 37 °C with secondary antibody diluted in 0.2% gelatin, washed three times in PBS-CM, and finally mounted using Fluoromount (Invitrogen).

### Cell biology image analysis and quantification

Fixed cells were imaged on a Carl Zeiss LSM 700 confocal laser scanning microscope running ZEN2011 software, using a Plan-Apochromat 40x/1.30 oil DIC objective and image settings were maintained the same for all images within one experiment. Images were acquired of cells that express similar levels of HA-KBP constructs based on immunostaining. Cells were selected on a first-come-first served basis. Images were processed and analysed using Fiji software^74^. For quantification of PEX transport, an ROI of the cell area was drawn and from this a second ROI at 5 - 7.5 μm from the outer cell area was created. A threshold ranging from 7500-10000 was determined for each experiment and for both ROIs the area with fluorescent intensity above threshold was determined in the RFP channel. From these values the percentage of cell area above threshold in the cell periphery was calculated. Results were normalized to the experimental condition without rapalog.

### Pull-down experiments and western blotting

HEK293T cells were cultured in 50/50 DMEM (Lonza)/Ham’s F10 (Lonza) medium supplemented with 10% FCS (Sigma) and 1% penicillin/streptomycin (Sigma). One day before transfection cells were diluted and plated into 6-well plates. Cells were co-transfected with pCl-Neo-BirA, HA-tagged constructs and bioGFP-tagged constructs using MaxPEI (Polysciences) in a ratio of 3/1 PEI/DNA, according to the manufacturer’s protocol. After 24-hours of expression, cells were washed in ice-cold PBS and lysed in lysis buffer (100 mM TrisHCl pH 7.5, 150 mM NaCl, 1% Triton X-100, protease inhibitors (Roche)) for 30 minutes on ice. Lysates were cleared by 30-minute centrifugation at 13.2 krpm at 4°C and supernatants were incubated with blocked (incubation for 30 minutes at RT in 50 mM Tris–HCl pH 7.5, 150 mM KCl, 0.2 μg/μl chicken egg albumin) Streptavidin Dynabeads M-280 (Invitrogen) for 1.5 hours at 4°C. Beads were then washed five times with washing buffer (100 mM Tris– HCl pH 7.5, 250 mM NaCl, 0.5% Triton X-100) and proteins were eluded from the beads by boiling for 10 minutes at 95°C in 2x DTT+sample buffer (20% glycerol, 4% SDS, 200 mM DTT, 100 mM Tris–HCl pH 6.8, bromophenol blue).

Protein samples were run on 10% SDS-PAGE gels and transferred to nitrocellulose membranes (Bio-Rad) by semi-dry blotting at 16V for 1 hour. Membranes were blocked by incubation in 3% bovine serum albumin (BSA) in PBST (PBS supplemented with 0.02% Tween20) for 1 hour at room temperature. This was followed by overnight incubation with primary antibodies in 3% BSA-PBST. Membranes were washed three times with PBST, incubated with secondary antibody in 3% BSA-PBST for 1 hour at room temperature, and washed three times with PBST. Membranes were scanned using an Odyssey Infrared Imaging system (LI-COR Biosciences) and blots were acquired at 680 nm and 800 nm.

## Acknowledgements

J.A. was supported by a grant from the Medical Research Council (MRC), U.K. (MR/R000352/1) to C.A.M, J.L. and A.P were supported by a grant from Worldwide Cancer Research, U.K. (16-0037) awarded to J.L and C.A.M. We thank Dr Alexander Cook for technical and processing assistance at the ISMB and Dr Radostin Danev at the Graduate School of Medicine, The University of Tokyo for custom processing scripts. Cryo-EM data collected at the Institute of Structural and Molecular Biology (ISMB), Birkbeck was on equipment funded by the Wellcome Trust, U.K. (202679/Z/16/Z, 206166/Z/17/Z and 079605/Z/06/Z) and the Biotechnology and Biological Sciences Research Council (BBSRC) UK (BB/L014211/1). We thank Dr Natasha Lukoyanova for support during data collection at the ISMB. For the remaining EM data collection, we acknowledge Diamond for access and support to the Electron Bioimaging Centre (eBIC) at Diamond, Harwell, UK, funded by the MRC, BBSRC and Wellcome Trust, U.K. S.S.R. was supported by a grant from the National Institute of General Medical Sciences (R01GM130556). N.O and M.O.S were supported by a grant awarded to M.O.S from the Swiss National Science Foundation (31003A_166608). J.J.A. was supported by the Netherlands Organization for Scientific Research (NWO-ALW-VICI, CCH), and the European Research Council (ERC) (ERC-consolidator, CCH).

## Author contributions

J.A, C.A.M, J.J.A.H and C.C.H performed experiments, analysed data and wrote the paper. C.A.M and C.C.H coordinated the project. J.A performed and analysed cryo-EM experiments on KBP and KBP-kinesin complexes, J.L and A.P performed cryo-EM experiments on Kif15_MD-MTs and J.A, J.L and A.P analysed these data. J.J.A.H performed and analysed inducible peroxisome motility and pull-down assays. N.O, S.R and M.O.S provided protein, DNA and/or background knowledge towards the experiments.

**The authors have no competing financial interests**

## Figure Legends

**Supplementary Figure 1.**
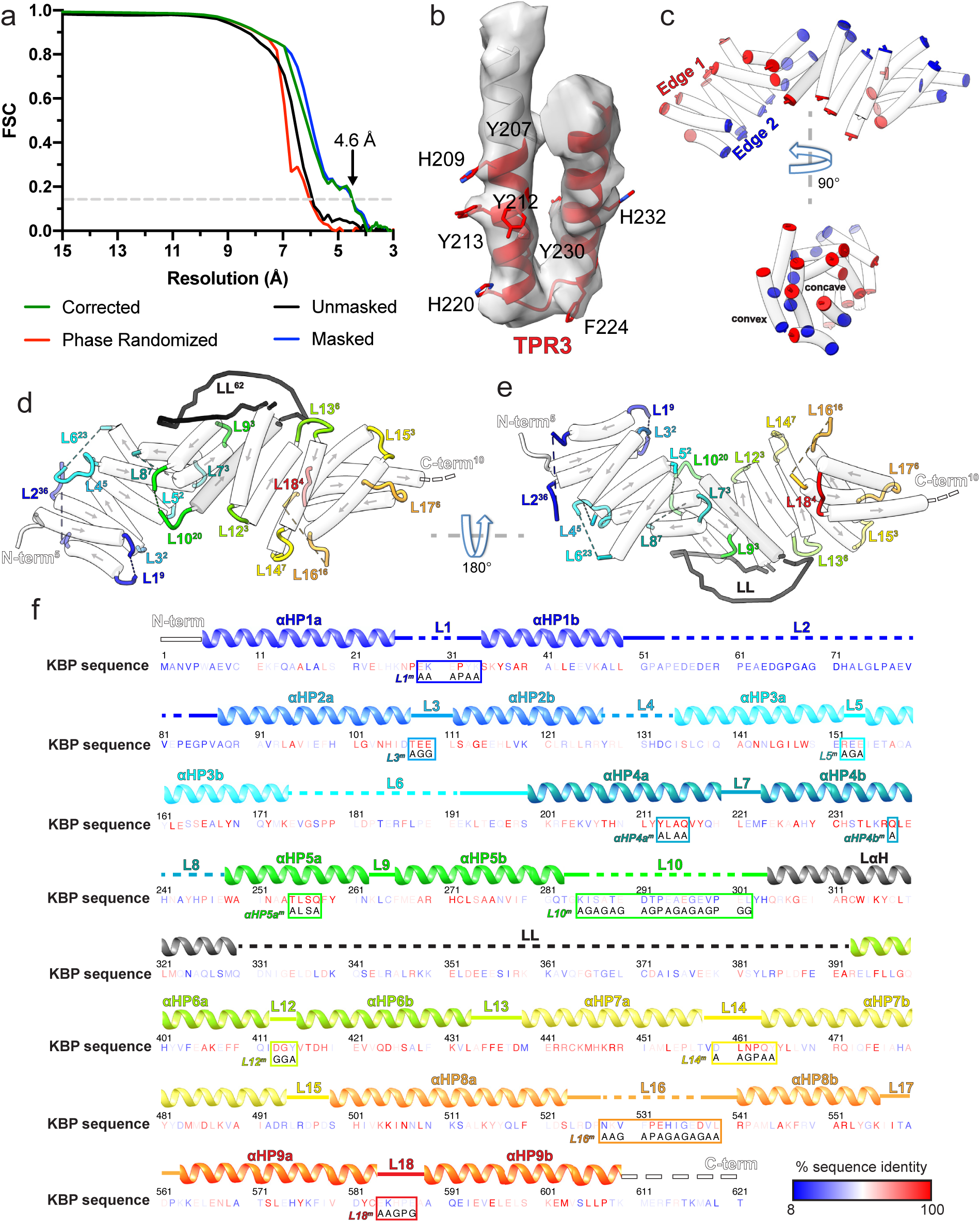
KBP reconstruction, loops, sequence, inter-species conservation and experimental mutations. (a) Gold-standard FSC curves between independent masked, unmasked, phase-randomised and corrected half-maps^76^ of KBP as calculated by RELION v3.1^51^ (4.6 Å resolution at the ‘gold-standard’ 0.143 FSC cutoff). (b) Density and fitted model for TPR3 of KBP, showing exemplar bulky side chain density that guided modelling. (c) Same view as Fig. 1c (upper panel), or rotated 90° around the indicated axis (lower panel), showing only α-helices (semi-transparent white tubes) with their terminal residues coloured, illustrating the edges and faces (concave and convex) of the α-solenoid respectively. (d) Same view as panel c, but now with loops shown and coloured (semi-transparent tube helices have their directionality represented by arrows). Each loop or terminus label has a superscript number indicating their length. (e) Same as panel d, rotated 180° around the indicated axis. (f) The human KBP sequence (numbering above), with residues coloured by intra-species sequence identity as indicated in the key. The following species were included in the Clustal Omega multiple sequence alignment^66^; *Homo sapiens, Mus musculus, Gallus gallus, Xenopus tropicalis, Alligator mississippiensis, Danio rerio, Drosophila melanogaster, Amphimedon queenslandica, Stylophora pistillata, Trichoplax adhaerens, Spizellomyces punctatus* and *Salpingoeca rosetta.* Above the sequence, secondary structure elements are indicated, coloured to delineate the nine α-helical pairs and connecting loops. Mutation sites are indicated within each boxed sequence region, labelled to coordinate with Fig. 5. Within these boxes, the mutated sequence is shown below the original wild-type sequence.

**Supplementary Figure 2.**
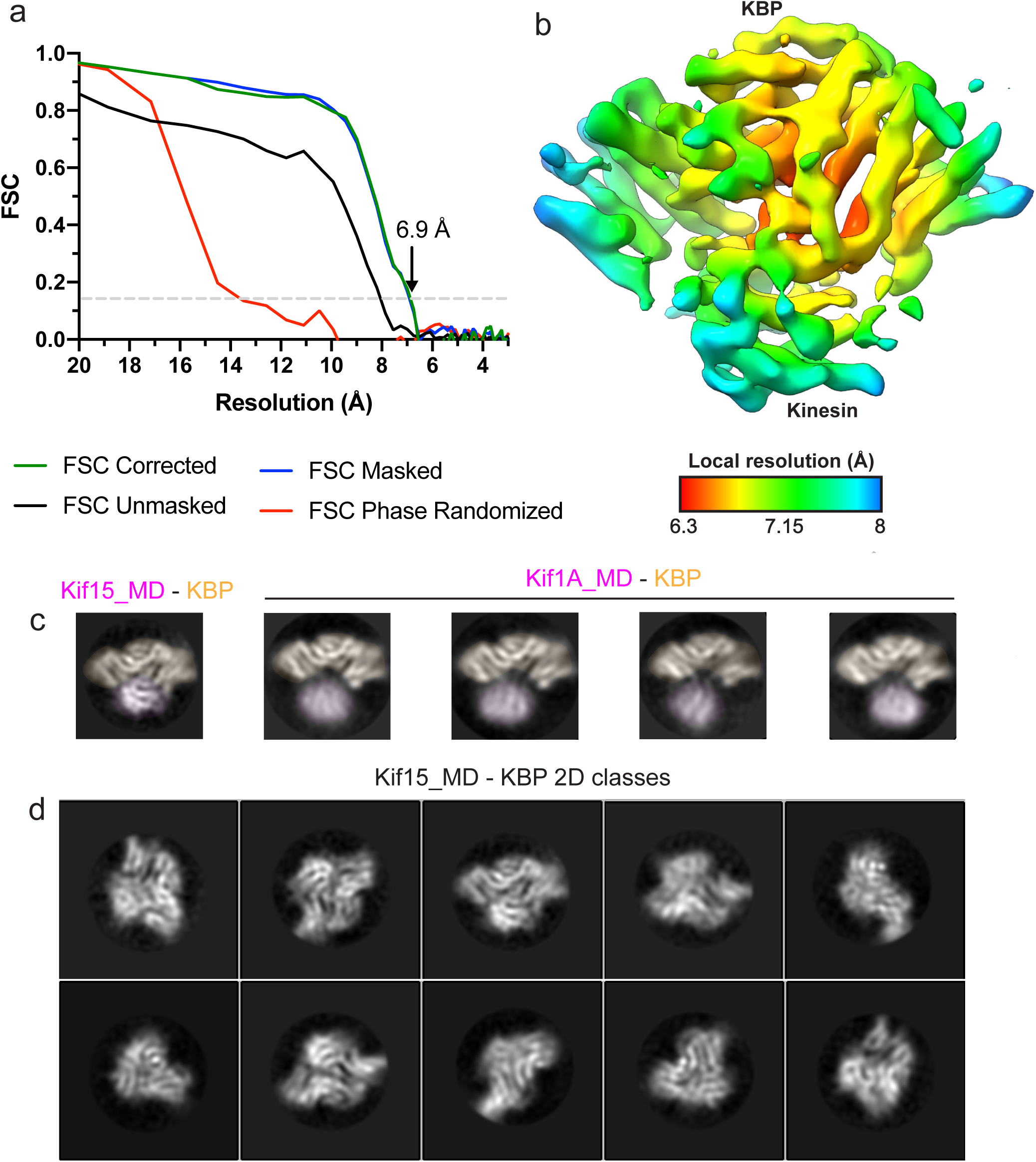
KBP-Kif15_MD reconstruction resolution estimation and 2D class analysis of Kif1A_MD-KBP and Kif15_MD-KBP complexes. (a) Gold-standard FSC curves between independent masked, unmasked, phase-randomised and corrected half-maps^76^ of the KBP-Kif15_MD complex as calculated by RELION v3.0^51^. The resolution at the ‘gold-standard’ 0.143 FSC cutoff is 6.9 Å. (b) Local resolution as calculated by RELION v3.0’s internal software, shown on the same view as in Fig. 2a with coloured density corresponding to the local resolutions indicated in the key. (c) Selected RELION v3.0^51^ 2D classes of KBP-Kif15_MD (left) and Kif1A_MD-KBP (4 to the right). Densities for the kinesin motor domain and KBP are pseudo-coloured pale magenta and pale orange respectively. Classes have been in-plane rotated such that KBP is seen from roughly the same orientation. Note poor resolution and a variable relative position in the Kif1A_MD. (d) A representative subset of Kif15_MD-KBP complex 2D classes, showing multiple orientations.

**Supplementary Figure 3.**
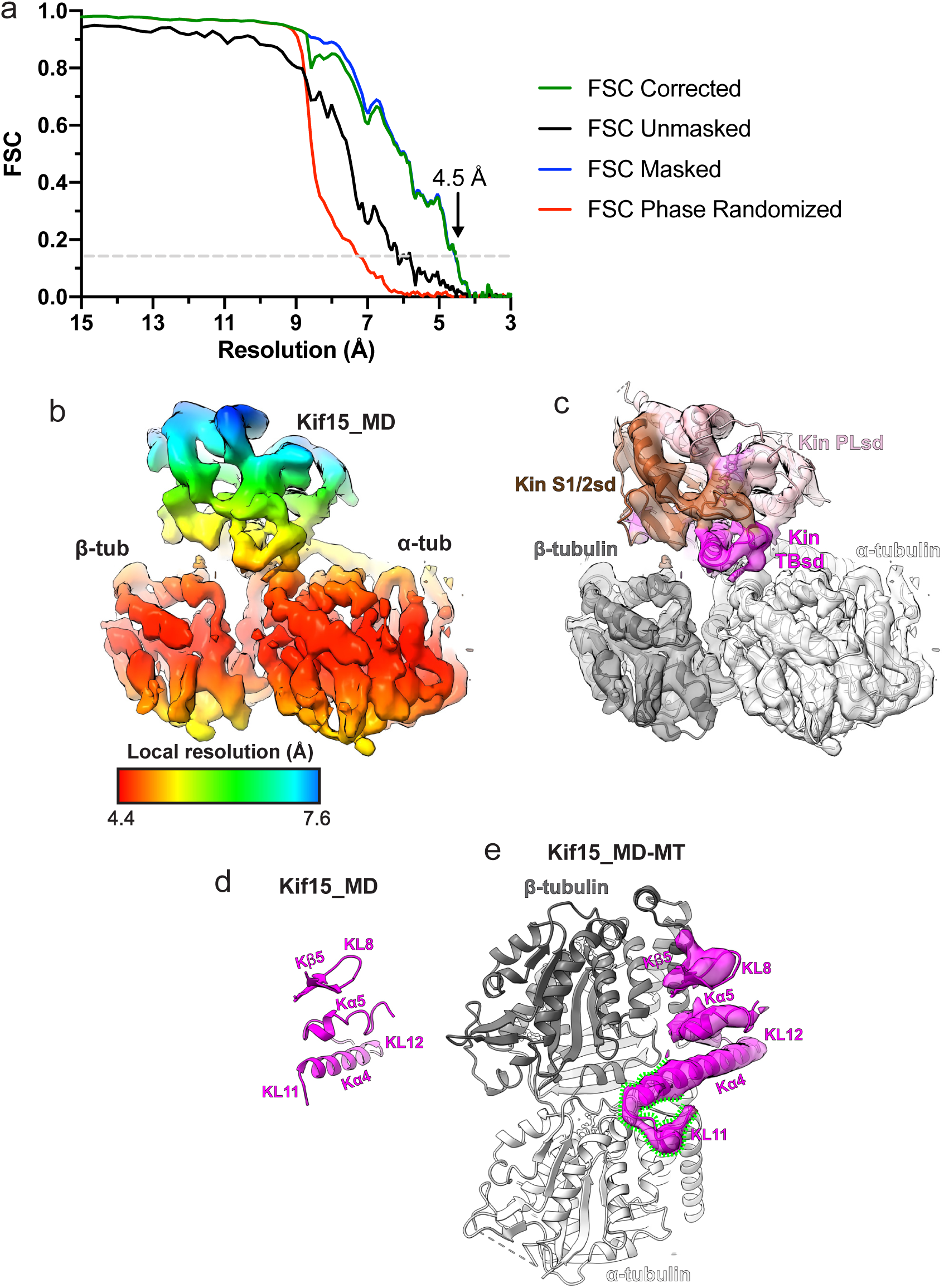
Kif15_MD adopts a canonical MT-bound kinesin conformation. (a) Gold-standard FSC curves between independent masked, unmasked, phase-randomised and corrected half-maps^76^ of the Kif15_MD-MT complex as calculated by RELION v3.0^51^. The resolution at the ‘gold-standard’ 0.143 FSC cutoff is 4.5 Å. (b) Local resolution as calculated by RELION v3.0’s internal software, with coloured density corresponding to the local resolutions indicated in the key. (c) The Kif15_MD-MT asymmetric unit model in corresponding density. The Kif15_MD is coloured by subdomain, bound Mg_2+_-AMPPNP coloured lilac and α and β-tubulin are coloured light and dark grey respectively, along with their corresponding cryo-EM densities. The same view as in panel b. (d) The tubulin binding subdomain of Kif15_MD alone from the crystal structure (PDB code:4BN2^28^). (e) Kif15_MD complexed with MTs with only the tubulin-binding subdomain and α and β-tubulin shown, along with the tubulin binding subdomain cryo-EM density (semi-transparent). Kα4 of Kif15_MD is extended relative to the crystal structure and KL12 adopts a new conformation on α-tubulin, as indicated by the green dashed lines. Panels d and e show the Kif15_MD tubulin-binding subdomain from the same viewpoint.

**Supplementary Figure 4.**
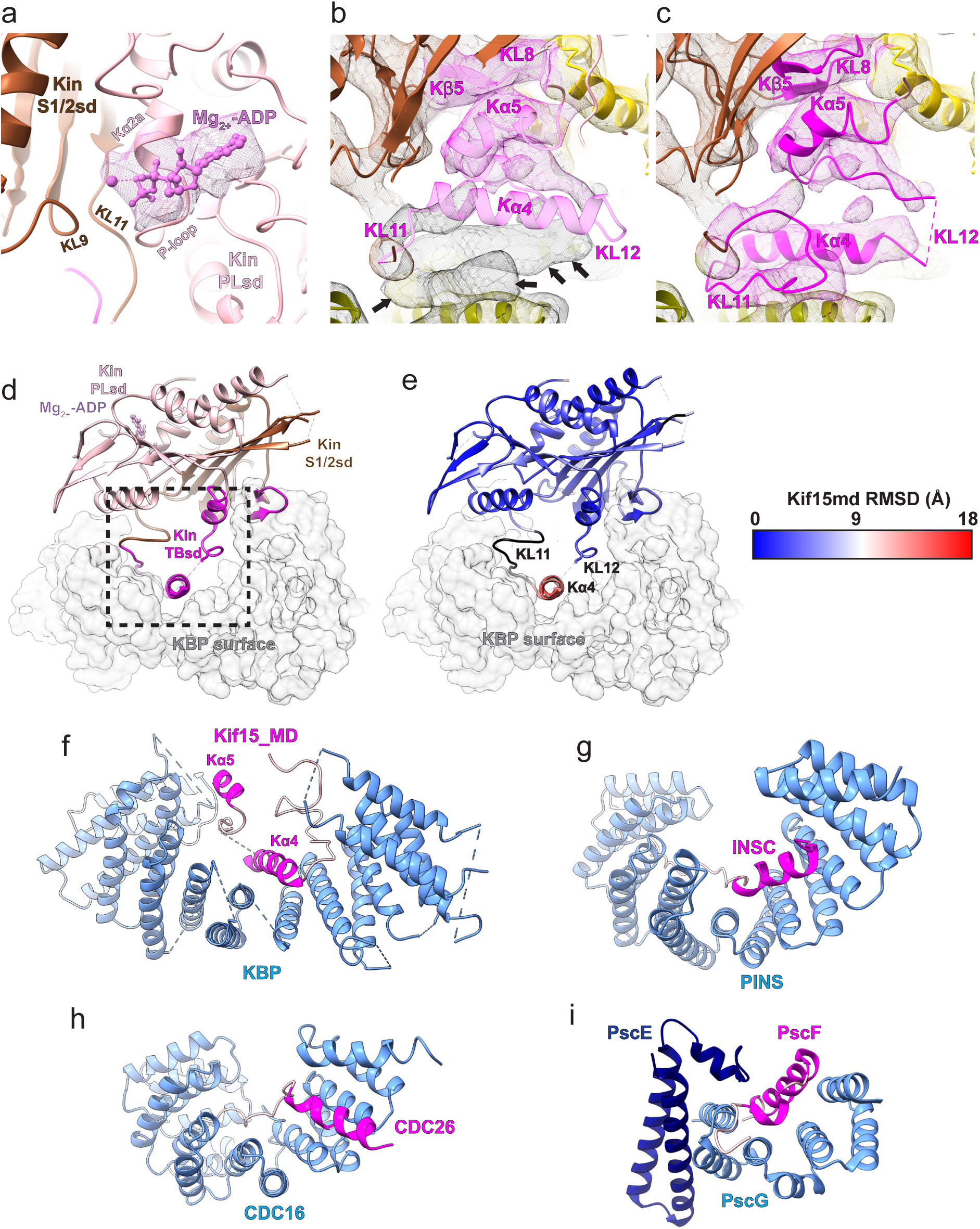
Movement of Kα4 of the Kin TBsd upon KBP binding and examples of TPR-containing α-solenoid proteins binding α-helical SSE ligands. (a) The Kif15_MD alone crystal structure (PDB code:4BN2^28^) is shown coloured by kinesin subdomain (as in Figure 3), fitted into the KBP-Kif15_MD complex cryo-EM map, with only density shown for the Mg_2+_-ADP as mesh. Density for Mg_2+_-ADP is found in the expected position between nucleotide binding elements KL9, KL11, Kα2a and the P-loop. (b) As in Figure 3a, but with a clipped viewpoint zoomed on the TBsd (in pale magenta to illustrate poor fit). Black arrows indicate unaccounted-for density. (c) As in Figure 3b, with the clipped viewpoint as in panel b of this figure (TBsd now opaque to illustrate good density fit). (d) The KBP-Kif15_MD model is shown as opaque ribbons, with kinesin subdomain colouring as in panel a and b and as labelled. KBP is shown as a transparent light grey surface representation. The boxed region indicates that shown in Fig. 3c-e. (e) RMSD in Å corresponding to the Kif15_MD overlay in panel d, shown on the model of Kif15_MD in complex with KBP (grey transparent surface). Parts of the model coloured black are disordered/missing in the Kif15_MD alone crystal structure. (f-i) Comparison of (f) KBP-Kif15_MD complex with other TPR-containing α-solenoids shown in blue, and binding peptide motifs shown in magenta or pink for helical and random coil regions respectively; (g) the PINS-INSC complex^30^, (h) the CDC16-CDC26 complex^31^ and (i) the PscE/PscG-PscF complex^32^.

**Supplementary Figure 5.**
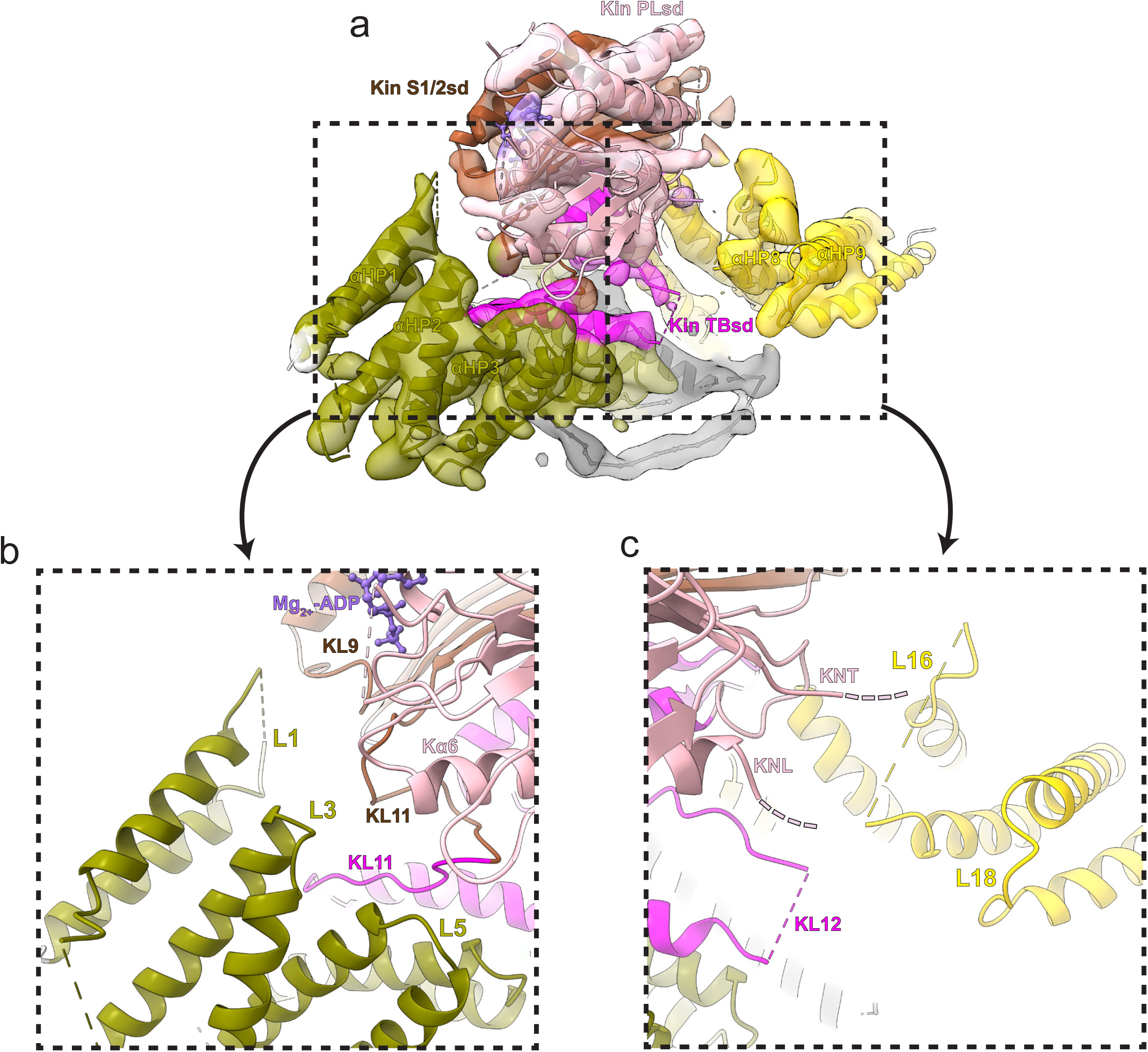
Additional KBP α-solenoid edge loops proximal to Kif15_MD. Colouring and representation as in Fig. 4. (a) A view showing α-helical pairs αHP1, αHP2, αHP3, αHP8 and αHP9 of KBP and the Kif15_MD coloured by subdomain as labelled. (b) Left zoomed region in panel a, with density removed, showing KBP L1, L3, L5 and proximal kinesin elements KL9, KL11 and Kα6. ADP is coloured in light orchid. (c) Right zoomed region in panel a, with density removed, showing KBP L16 and L18 and proximal Kif15_MD elements KNT (kinesin N-terminus), KNL (kinesin neck-linker) and KL12.

**Supplementary Figure 6.**
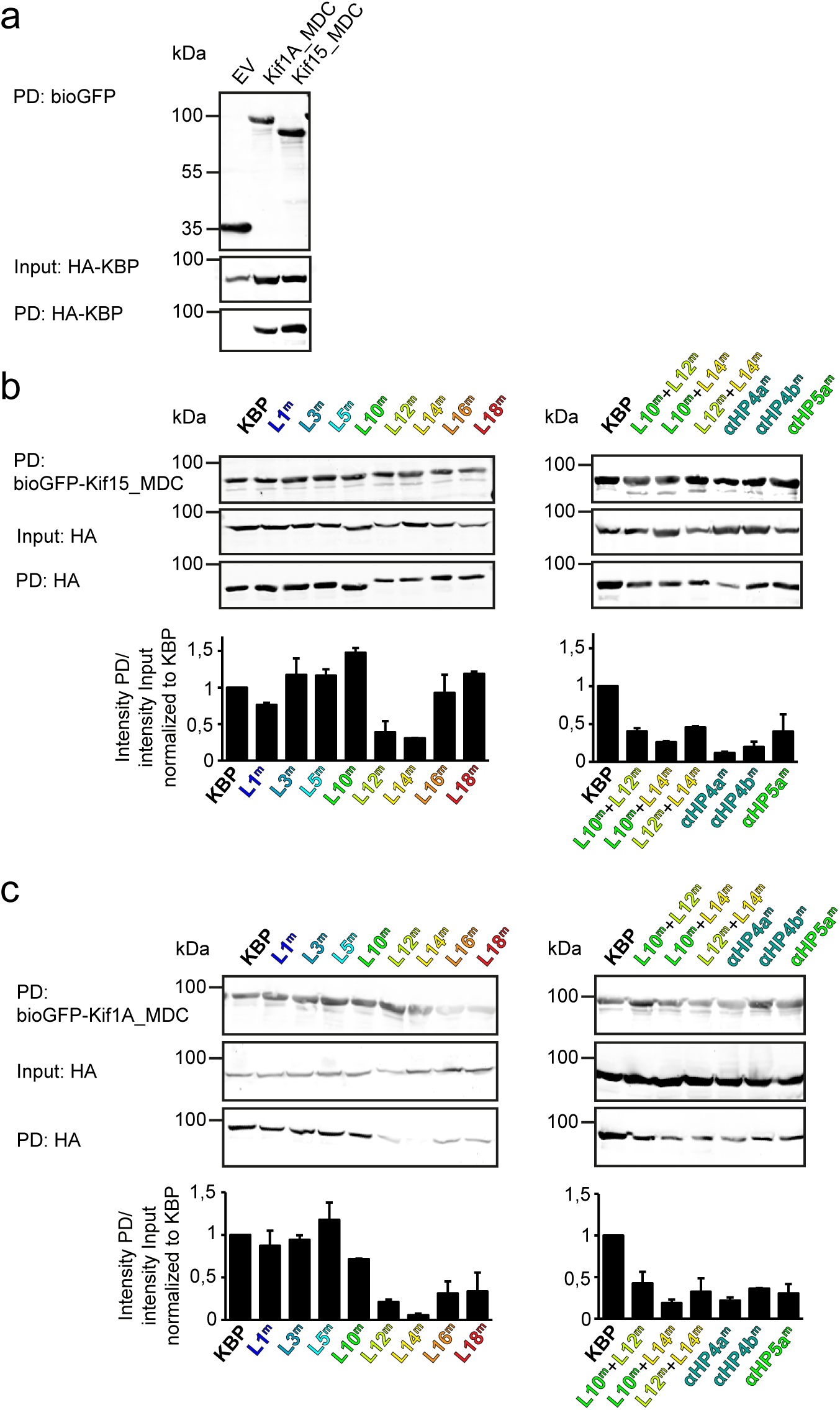
Pull-down experiments demonstrate the effect of KBP mutation on the interaction between Kif15 and Kif1A. (a) Control pull-down experiment with bioGFP-EV, bioGFP-Kif1A_MDC or bioGFP-Kif15_MDC and HA-KBP showing that KBP interacts with Kif1A_MDC and Kif15_MDC, but not with bioGFP-EV. (b, c) Example of pull-down experiment showing the interaction between (b) Kif15_MDC or (c) Kif1A_MDC and mutated KBP constructs in HEK293T cell lysates. Graphs underneath show the quantification of the intensity of the mutated HA-KBP construct in the pull-down fraction over the input fraction and normalized to HA-KBP. Data are displayed as mean ± s.e.m. (data from two independent experiments).

**Supplementary Video 1. KBP undergoes conformational change to relieve clashes when forming a complex with Kif15_MD.**

The KBP-alone model was superimposed on the KBP-Kif15_MD model using UCSF Chimera’s matchmaker^56^. A conformational morph movie was then generated in Chimera between the KBP-alone and Kif15_MD bound states, with Kif15_MD shown throughout to illustrate the relief of clashes. The N-terminal and C-terminal subdomains are coloured in olive and gold respectively, as in Fig. 2a,b, while Kif15_MD is shown in pale magenta. Distances between identified clashing atoms when KBP-alone is superimposed onto the KBP-Kif15_MD model are indicated by red linking lines and KBP clashing residues and side chains shown in cyan. Atoms that were clashing remain coloured while the red lines gradually disappear as the clashes are relieved by the conformational change. Clashes were calculated in Chimera using default criteria.

**Supplementary Video 2. Interaction of KBP with the Kif15 MD.**

Model of the KBP-Kif15_MD complex (ribbon representation) displayed in experimental cryo-EM density. The N-terminal (olive) and C-terminal (gold) subdomains and the linker region (black) are shown in KBP, while the Kif15_MD Switch 1/2 subdomain (Switch 1/2 subdomain) is coloured sienna, the P-loop subdomain (Kin-PLsd) is coloured light pink and the Kif15_MD tubulin-binding subdomain (TBsd) is coloured magenta. Semi-transparent density is coloured regionally as per the fitted model.

**Supplementary Table 1.** Cryo-EM reconstruction information and model refinement statistics and model geometry. Data collection, processing and model refinement information for the KBP, KBP-Kif15_MD and Kif15_MD-MT datasets.

|  | **KBP**<br>**(EMDB: EMD-11338, PDB: 6ZPG)** | **KBP-Kif15_MD**<br>**(EMDB: EMD-11339, PDB: 6ZPH)** | **Kif15_MD-MT**<br>**(EMDB: EMD-11340, PDB: 6ZPI)** |
| --- | --- | --- | --- |
| **Data collection and processing** |  |  |  |
| Pixel size (Å)* | 1.055, 1.043 or 1.047 | 1.047 | 1.39 |
| Number of micrographs (collected, final)* | 9360, 7547 | 6497, 5138 | 214,202 |
| Final particle number | 258,049 | 7,513 | 12,674 |
| Map resolution (Å) | 4.6 | 6.9 | 4.5 |
| FSC threshold† | Independent half-map FSC 0.143 | Independent half-map FSC 0.143 | Independent half-map FSC 0.143 |
| **Refinement** |  |  |  |
| Refinement resolution (Å) | **4.6** | **6.9** | **6** |
| CC_mask‡ | **0.64** | **0.74** | **0.60** |
| Map sharpening *B*-factor (Å<sup>2</sup>) | **-200** | **-495** | **-134** |
| Model composition |  |  |  |
| Nonhydrogen atoms | **3,808** | **6,232** | **9,420** |
| Protein residues | **610** | **948** | **1185** |
| Ligands | **0** | **1** | **4** |
| R.m.s. deviations§ |  |  |  |
| Bond lengths (Å) | **0.01** | **0.01** | **0.08** |
| Bond angles (°) | **0.96** | **1.07** | **0.17** |
| Validation# |  |  |  |
| MolProbity score | **1.66** | **1.84** | **1.95** |
| Clashscore | **5.25** | **7.31** | **13.25** |
| Poor rotamers (%) | **0.5%** | **0.9%** | **0.1%** |
| Ramachandran plot# |  |  |  |
| Favored (%) | **94.38** | **93.13** | **95.38** |
| Allowed (%) | **5.62** | **6.87** | **4.62** |
| Outliers (%) | **0** | **0** | **0** |

\*Inclusive of all data collection sessions
†The resolution value at the gold-standard Fourier Shell Correlation (FSC) 0.143 criterion between independently refined half-maps.
‡Cross-correlation provided by Phenix real-space refine<sup>7</sup>.
§Root-mean-square deviations of bond lengths or angles in the model.
#As defined by the MolProbity validation server<sup>77</sup>.

**Supplementary Table 2.**
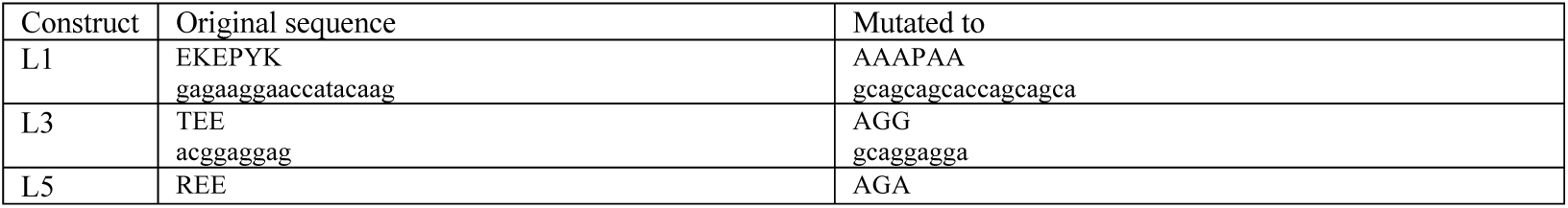
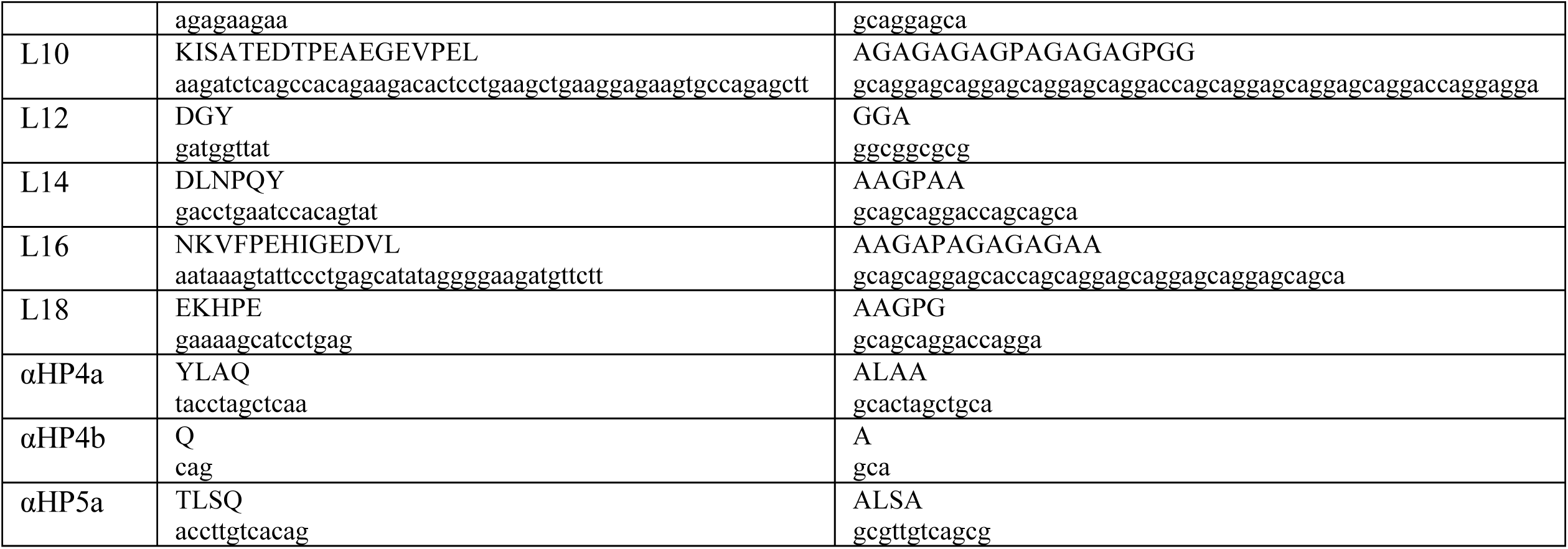
KBP mutants used in this study. The original and mutated amino acid (top) and nucleotide sequences (bottom) are shown for each construct.

